# Do naps benefit novel word learning? Developmental differences and white matter correlates

**DOI:** 10.1101/2021.11.22.469237

**Authors:** E. van Rijn, A. Gouws, S. Walker, V. C. P. Knowland, S. A. Cairney, M. G. Gaskell, L. M. Henderson

## Abstract

Behavioural and neuroimaging data suggest that memory representations of newly learned words undergo changes during nocturnal sleep, including improvements in explicit recall and lexical integration (i.e., after sleep, novel words compete with existing words during online word recognition). Some studies have revealed larger sleep-benefits in children relative to adults. However, whether daytime naps play a similar facilitatory role is unclear. We investigated the effect of a daytime nap (relative to wake) on explicit memory (recall/recognition) and lexical integration (lexical competition) of newly learned novel words in young adults and children aged 10-12 years, also exploring white matter correlates of the pre- and post-nap effects of word learning in the child group with diffusion weighted MRI. In both age groups, a nap maintained explicit memory of novel words and wake led to forgetting. However, there was an age group interaction when comparing change in recall over the nap: children showed a slight improvement whereas adults showed a slight decline. There was no evidence of lexical integration at any point. Although children spent proportionally more time in slow-wave sleep (SWS) than adults, neither SWS nor spindle parameters correlated with over-nap changes in word learning. For children, increased fractional anisotropy (FA) in the uncinate fasciculus and arcuate fasciculus were associated with the recognition of novel words immediately after learning, and FA in the right arcuate fasciculus was further associated with changes in recall of novel words over a nap, supporting the importance of these tracts in the word learning and consolidation process. These findings point to a protective role of naps in word learning, and emphasize the need to advance theories of word learning by better understanding both the active and passive roles that sleep plays in supporting vocabulary consolidation over development.

## 1. Introduction

Vocabulary knowledge is not only a critical component of language and communication, it is a longitudinal predictor of an array of life outcomes, from school grades to employment and mental and physical health (Armstrong et al., 2017). Understanding the underlying neurocognitive mechanisms of word learning is valuable for advancing theory, and can also inform how to promote vocabulary learning over the lifespan. Numerous studies have shown that the journey of a new word, from unfamiliar to established, plays out over extended period, with sleep actively supporting long-term retention (James et al., 2017; Kurdziel et al., 2017). Nocturnal sleep has been found to benefit word learning in both children (Brown et al., 2012; Henderson et al., 2012) and adults (Dumay & Gaskell, 2007; Tamminen et al., 2010), and in some studies these benefits have been larger in children as compared to adults (James et al., 2017). While evidence points to daytime naps also supporting word learning in adults (e.g., Tamminen et al., 2017), it is less clear if this is also the case for children. Furthermore, little is known about the neurocognitive and neuroanatomical mechanisms that underlie new word learning over a nap, and crucially, whether naps (like nocturnal sleep) might be more beneficial for younger than adult word learners. These are important questions, especially given calls to incorporate naps into the school day to improve learning outcomes (Cabral et al., 2018; Ji et al., 2019; Lemos et al., 2014).

### 1.1. Sleep and word learning

Word learning is a key example of declarative memory consolidation, where sleep plays a well-established role (see Rasch & Born, 2013, for a review). Numerous studies have shown that nocturnal sleep (relative to equivalent periods of wake) can strengthen explicit memories of newly learned words (i.e., leading to gains in recall and recognition) and allow them to become integrated in long-term lexical memory in both adults (Dumay & Gaskell, 2007; Tamminen et al., 2010) and children (Henderson et al., 2012; James et al., 2020; Smith et al., 2018). Lexical integration can be measured via the degree to which spoken novel non-words (e.g., “dolpheg”) engage in lexical competition with words that have overlapping phonology (e.g. “dolphin”), a process that is fundamental to the recognition of established lexical entries (Gaskell & Dumay, 2003). For example, using the pause detection task (Mattys & Clark, 2002), Dumay and Gaskell (2007) showed that adults were slower to make speeded decisions about the presence or absence of a silent pause inserted into existing, familiar words (e.g., “dolph_in”) that overlapped with newly trained novel words (e.g., “dolpheg”) than familiar words that did not overlap with newly trained words. Assuming that the new competitors had become integrated in the language network, the longer latencies for existing words which now had these new competitors were argued to reflect increased lexical activity at pause onset which worked to decrease resources available for making the response. Importantly, however, this “lexical competition” effect did not emerge until after a period of sleep (see Henderson et al., 2012, for similar findings with children; although see Kapnoula et al., 2015; Lindsay & Gaskell, 2013, for examples of where lexical competition can emerge without sleep).

This pattern of word learning has been explained in terms of a complementary learning systems (CLS) account (Davis & Gaskell, 2009). Classic CLS models of declarative memory propose a dual memory system, with the hippocampus serving as a short-term store for new memories and the neocortex as a site for long-term storage (McClelland et al., 1995). As applied to word learning, the initial retrieval of word meaning and form requires hippocampal mediation, but after a period of consolidation, an integrated neocortical representation is strengthened. This prolonged process of consolidation has been suggested to occur offline, and specifically during sleep (Davis et al., 2009; Davis & Gaskell, 2009; Dumay & Gaskell, 2007).

The CLS account of word learning is supported by functional magnetic resonance imaging (fMRI) findings. In adults, Davis et al. (2009) observed increased neural responses to untrained pseudowords and pseudowords trained on the day of an fMRI scan relative to pseudowords trained on the day before the scan. This response difference was seen in brain regions that support semantic access and effortful phonological processing (e.g., in the bilateral superior temporal gyrus, the inferior frontal, premotor and motor cortices, and in the cerebellum). In keeping with the CLS account, the authors argued that sleep worked to improve the efficiency with which new words are represented in cortical systems involved in word recognition and production. Davis et al. (2009) also observed greater hippocampal activity for the untrained pseudowords, with this hippocampal involvement decreasing over the night after learning.

Building on this, Landi et al. (2018) trained a slightly younger sample (16-25 years) on associations between spoken pseudowords and pictures of rare minerals or fish, half were learned in the evening (consolidated trained words) and half the next morning before scanning (unconsolidated trained words). Consolidated, compared to unconsolidated, trained words elicited increased activation in language regions, including the left superior and medial temporal gyrus, left inferior parietal lobule, posterior cingulate and precuneus. However, unlike Davis et al. (2009) where hippocampal responses decreased for consolidated words, Landi et al found that hippocampal activity was greater for all trained words (consolidated and unconsolidated) relative to novel untrained pseudowords. Whilst this difference in the hippocampal response could be attributed to the slightly younger sample used in Landi et al, it may also be due to the more complex stimuli that were trained, which may have remained hippocampally-dependent even after overnight consolidation.

Studying younger children and adolescents, Takashima et al. (2019) measured behavioural and fMRI responses to newly learned Japanese words in 8-10 and 14-16 year olds, both immediately after training and one week later. Aligning with Davis et al. (2009), hippocampal involvement decreased with time; however, in contrast, there was no evidence of cortical involvement even one week after training. Additionally, while Davis et al. (2009) found behavioural evidence for lexical integration of novel words the day after learning in adults, Takashima et al. (2019) did not find this one week later. This suggests that for children, novel words may undergo a more protracted phase of lexical integration than in adults. Interestingly, this differs from the conclusions of studies that have used purely behavioural measures, where there are claims of sleep-dependent lexical integration as little as 12-hours after training (e.g., Henderson et al., 2012). Together, these findings highlight the value of using both behavioural and neuroimaging measures in parallel when attempting to understand the neurocognitive mechanisms that support word learning, particularly when examining developmental differences. The developmental differences in hippocampal involvement in the above studies also raises the possibility that the original proposal of declining hippocampal involvement after sleep (i.e., Davis et al., 2009) may be overly simplistic. Indeed, studies of rapid cortical plasticity in word learning in adults (e.g., Hofstetter et al., 2017; Shtyrov et al., 2010; Vukovic et al., 2021) point to a structural basis for rapid word encoding mechanisms, highlighting the importance of studying the baseline integrity of white matter structures involved in language acquisition.

Building on the classic CLS model, the *Active Systems* hypothesis provides a mechanistic explanation for *how* sleep supports systems consolidation (Born & Wilhelm, 2012; Diekelmann & Born, 2010; Rasch & Born, 2013). During non-rapid eye movement (NREM) sleep, and slow wave sleep (SWS) in particular, newly encoded hippocampal memories are argued to be repeatedly reactivated until they are redistributed to the neocortex, where they are strengthened and integrated within pre-existing neural networks. This process of redistribution relies on the interplay between three key brain oscillations during sleep: neocortical slow oscillations, thalamo-cortical sleep spindles and hippocampal sharp-wave ripples (Born & Wilhelm, 2012; Diekelmann & Born, 2010; Rasch & Born, 2013). NREM sleep spindles and characteristics of SWS, including delta power (1-4 Hz), slow wave activity (SWA, 0.5 - 4 Hz) and time spent in SWS, have been linked to declarative memory consolidation (Diekelmann et al., 2012; Marshall & Born, 2007; Mölle et al., 2004; Plihal & Born, 1997). Sleep spindles consist of short bursts of oscillatory activity in the sigma frequency band (De Gennaro & Ferrara, 2003; Muehlroth et al., 2019) and can be categorized into frontal slow spindles (9-12.5 Hz) and central fast spindles (12.5-16 Hz) (Muehlroth et al., 2019). Behavioural measures of sleep-dependent memory consolidation have repeatedly been associated with fast spindle density (Barakat et al., 2011; Chatburn et al., 2013; Hahn et al., 2019; Tamaki et al., 2008) and power within the fast spindle frequency band (Cairney et al., 2018; Mölle et al., 2011).

Sleep-related changes in new word knowledge have also been associated with sleep spindle and slow-wave parameters in adults (Tamminen et al., 2013; Tamminen et al., 2010; Weighall et al., 2017) and children (Fletcher et al., 2020; Smith et al., 2018). Tamminen et al. (2010) found significant correlations between overnight changes in lexical competition for both slow and fast spindle types in adults. Additionally, more time in SWS was associated with larger decreases in recognition reaction time. Weighall et al. (2017) found that explicit memory for novel words after sleep correlated with fast (but not slow) spindle density in adults. In typically developing children aged 8-13 years, Smith et al. (2018) found that overnight change lexical competition correlated positively with spindle power averaged over fast and slow frequency ranges. Overnight improvements in explicit memory (i.e., captured by a cued recall task) were also related to spindle power, as well as slow wave activity (SWA). Extant evidence from polysomnography studies therefore suggests that sleep supports the consolidation of newly learned words similarly in childhood and adulthood. However, sleep architecture undergoes significant changes across the lifespan (Ohayon et al., 2004), with these changes potentially leading to subtle developmental differences in consolidation mechanisms (James et al., 2017). Amongst these architectural changes are increases in the amount of nocturnal SWS as well as higher SWA over childhood, with claims that the proportion of SWS peaks at age 10-12 years (Campbell & Feinberg, 2009; Kurth et al., 2010; Ohayon et al., 2004; Wilhelm et al., 2012; Wilhelm et al., 2013). Since SWS is thought to be key for effective sleep-dependent memory consolidation and tends to account for roughly double the proportion of night time sleep in pre-early adolescents relative to adults, it has been predicted that direct comparisons of these age groups will produce enhanced sleep-dependent consolidation effects in children. Aligning with this, Wilhelm et al. (2013) found that compared to adults, 8-11 year old children showed greater gains in explicit knowledge of a motor sequence following sleep, and this was related to higher nocturnal SWA (see also Peiffer et al., 2020). For word learning, Weighall et al. (2017) demonstrated that after sleep, 7-8 year-olds had greater increases in explicit memory for newly learned words than adults. James et al. (2019) also found that children (7-9 years old) showed bigger overnight improvements in recall of novel words 24 hours after initial learning than compared to adults, and they continued to improve to a greater extent when tested one week later. Bishop, Barry & Hardiman (2012) also found that children outperformed adults on the offline retention of novel word forms, using a nonword repetition task, even when initial performance was matched. However, since James et al. (2019) and Bishop et al (2012) did not measure sleep directly and Weighall et al. (2017) did not measure sleep in both adults and children, these enhanced effects in children cannot be specifically attributed to greater amounts of SWS. Furthermore, enhanced consolidation effects in children are difficult to reconcile with Takashima et al. (2019), who observed no evidence of increased cortical responses to newly learned words after sleep in children. This raises the possibility that the enhanced explicit memory benefits previously found in children are not purely a consequence of enhanced neocortical consolidation over sleep.

### 1.2. Daytime naps and word learning

Naps, like nocturnal sleep, have been shown to promote active systems consolidation for newly acquired declarative memories (e.g., Cairney et al., 2018; Farhadian et al., 2021; Kurdziel et al., 2013; Lokhandwala & Spencer, 2021; Tucker & Fishbein, 2008). However, the effect of naps on word learning in adults or children has not been studied extensively. For adults, there is preliminary evidence that naps play a more protective role in word learning (Tamminen et al., 2017), but it is unclear if this is the same for children.

Studying adults, Tamminen et al. (2017) found that a 90-minute daytime nap protected memories of newly learned novel words from forgetting on a free recall task, while an equal period of wake did not. However, over-nap change in free recall did not correlate with time spent in any sleep stage or with sleep spindle density. This differs from the strengthening or enhancing effects of nocturnal sleep on explicit memory for new words where changes over sleep *are* associated with slow wave and sleep spindle parameters (e.g., Dumay & Gaskell, 2007; Tamminen et al., 2010), potentially suggesting a passive role of nap-based sleep (e.g., offering protection from interference) as opposed to allowing for active systems consolidation. Furthermore, lexical integration effects were found following a delay regardless of whether it was filled with a nap or wake, which again differs from previous overnight studies in which lexical integration was only found after sleep (e.g., Dumay & Gaskell, 2007). However, Tamminen et al. (2017) used a lexical decision task to capture lexical integration, in which participants made timed overt judgements about lexicality. The pause detection task used by Dumay and Gaskell (2007) arguably captures a more implicit form of lexical competition that is less prone to strategic effects (i.e., noticing the overlap between novel and existing competitors during the task). Thus, whether online lexical competition emerges after a nap in adults, as it does after nocturnal sleep, remains unclear. It should also be noted that Tamminen et al. (2017) used a targeted memory reactivation paradigm, in which half of the trained words were “cued” (i.e., played through loudspeakers during SWS while participants remained asleep), thus any effects of the nap may have been influenced by this cueing. Indeed, they observed an association between change in lexical competition over the nap for cued words and time spent in REM sleep, but this was not observed for non-cued words. This again implies that (without cueing) naps might not play an active role in the consolidation of newly learned words.

Evidence broadly points to beneficial effects of napping on memory in pre-schoolers and young children (Giganti et al., 2014; Kurdziel et al., 2013). For instance, in 3-6-year-olds, memory of a storybook was better following a nap compared to an equal time of wake and change in post-nap performance was associated with time spent in SWS (Lokhandwala & Spencer, 2021). In adolescents, Piosczyk et al. (2013) reported that a nap can promote the retention of word-pairs in 16-year-old females (selected to reduce variance related to gender) (see also Lau et al., 2018), but only when the nap was characterised by high sigma power. However, there is a paucity of studies of novel word learning and lexical integration over a nap in children. It is therefore currently unknown whether the higher levels of SWS (and purported associated enhanced benefits of sleep on memory) characterise nap sleep in children relative to adults. In adults, daytime naps are reported to have significantly lower spindle density than night-time sleep (van Schalkwijk et al., 2019), thus it is plausible that the benefits of a nap for word learning may be reduced for both age groups, but this does not perturb the potential for developmental differences.

### 1.3. White matter correlates of word learning

As discussed, fMRI studies with adults have revealed, broadly speaking, that novel word representations initially engage the hippocampal system (Breitenstein et al., 2005; Davis et al., 2009). However, over time (and sleep), hippocampal activation declines and activation in cortical structures associated with the representation of words (e.g., left middle temporal cortex) persists or increases (Bakker-Marshall et al., 2018; Takashima et al., 2019; Landi et al., 2018). Yet, findings from Takashima et al. (2019) in 8-10 and 14-16 year old children suggest decreased hippocampal involvement in response to newly learned words one week after learning *without* corresponding increases in cortical activation. One possibility is that the structural properties of the developing language network may dictate how efficiently new words are strengthened in cortical networks.

In parallel with the growth of cortical grey matter, axonal bundles become increasingly myelinated over childhood and adolescence (Miller et al., 2012), which is crucial for the interneuronal communication that underlies cognitive development (Casey et al., 2005; Khundrakpam et al., 2013; Schmithorst et al., 2005). Diffusion weighted imaging (DWI), a non-invasive technique used to measure water diffusion characteristics as an index of the integrity of white matter microstructure, has consistently found non-uniform changes in white matter integrity across childhood and adolescence (see Lebel, Treit, et al., 2019, for a review). These studies typically measure fractional anisotropy (FA, the directional preference of diffusion), axial diffusivity (AD, diffusion along the main axis of diffusion), radial diffusivity (RD, rate of diffusion in the transverse direction) and mean diffusivity (MD, mean diffusion rate) (Soares et al., 2013). Various measures of verbal ability are associated with developmental changes in these metrics (Farah et al., 2020; Houston et al., 2019). For instance, greater conversational experience with adults at age 4-6 years is associated with stronger white matter connectivity (as reflected by a positive correlation with FA) in the left arcuate fasciculus (AF) and left superior longitudinal fasciculus (SLF) (Romeo et al., 2018; see also Su et al., 2018; Urger et al., 2015). The SLF and AF connect the frontal cortex to the occipital, parietal and temporal cortices and play a key role in speech and language processing in both the left and right hemispheres (Catani & Thiebaut de Schotten, 2008). Furthermore, auditory recall of word lists in children aged 9-15 years is associated with white matter integrity in the uncinate fasciculus (UF) (Mabbott et al., 2009; see also Lebel, Benischek, et al., 2019). The UF is the major white matter tract connecting frontal regions with the temporal pole, anterior temporal cortex, parahippocampal gyrus and amygdala (Von Der Heide et al., 2013), with communication between these cortical regions thought to underlie declarative memory skills (Schmahmann et al., 2007; Squire & Zola-Morgan, 1991) and semantic processing (Holland & Lambon Ralph, 2010; Papagno et al., 2011). Whilst associations between these tracts and language measures have predominantly been identified as left lateralized (Mabbott et al., 2009; Romeo et al., 2018; Sreedharan et al., 2015; Su et al., 2018; Urger et al., 2015), other studies report associations in the right (Farah et al., 2020) or both hemispheres (Romeo et al., 2018; Sreedharan et al., 2015).

To our knowledge, only one study has used DWI to examine associations of the microstructural properties of cortical regions and white matter tracts with novel word learning. Hofstetter et al. (2017) examined learning-induced changes in cortical plasticity following a one-hour training phase in which adults learned novel word-picture pairs. They found that learning of new lexical items induced immediate changes, including reductions in mean diffusivity, in a number of cortical regions linked to language (e.g., the inferior frontal gyrus, the middle temporal gyrus and the inferior parietal lobule). Of interest, they found that changes in diffusivity in the SLF were correlated with the rate at which adults learned the new words. However, Hofstetter et al. (2017) only examined the immediate consequences of the word learning process. To our knowledge, the white matter correlates of both immediate and delayed consequences of novel word learning (particularly in children) are yet to be examined.

### 1.4. Current study

This study focuses on three key questions. First, while nocturnal sleep benefits novel word learning and integration of novel words in the lexical network in adults and children, is the same true for naps? Second, do children benefit more from naps for novel word learning than adults? Third, what are the white matter correlates of novel word learning in children? We took a multi-method approach to addressing these questions, examining the neurocognitive (Experiments 1 and 2) and neuroanatomical (Experiment 3) mechanisms of word learning over a nap, in children and adults. Experiment 1 investigated whether a daytime nap (relative to an equivalent period of wake) led to benefits for explicit memory and lexical integration of newly learned novel words in young adults. Experiment 2 repeated this endeavour in children aged 10-12 years old, targeting the age at which SWS is generally in abundance (i.e., a larger proportion of time is spent in SWS and SWA shows greater amplitude) relative to adults (e.g., Wilhelm et al., 2013). Both of these experiments captured sleep architecture in the laboratory using polysomnography to examine whether over-nap changes in novel word memory were associated with electrophysiological parameters previously linked to active systems consolidation (i.e., spindle density, sigma power, delta power and time in SWS). A cross-experiment analysis explored developmental differences in nap architecture and the effect of napping on word learning. Finally, Experiment 3 used DWI to explore how the structural properties of the developing language system in the brain are associated with word learning, measured both immediately after learning and through capturing changes in memory over a nap.

## 2. Experiment 1: Daytime naps and novel word learning in adults

The following pre-registered hypotheses were tested (https://osf.io/63p2e):

1. There will be improvements in explicit memory for the novel words after a nap, but not after wake.
2. There will be a greater increase in lexical competition (i.e., slower responses to words for which a novel competitor has been trained, versus untrained words) following a nap compared to an equal period of wake.
3. The size of these sleep-based improvements in lexical integration and explicit memory will be positively associated with spindle density, sigma power^1^, delta power and time in SWS during the nap.

### 2.1. Material and methods

#### 2.1.1. Participants

Participants were all students aged 18-24 years at the University of York, who took part in two conditions, spaced approximately one week apart. For both weeks, participants were instructed to sleep normally and to not consume alcohol the night before testing. On the morning of testing, they were required to get up by 7am and to not nap or consume any caffeinated products. Actigraphy (Actiwatch Spectrum Plus, Philips Respironics Inc., Pittsburgh, Pennsylvania) was used to verify participants’ sleep schedule the night before the experiment. In total, 36 students took part. All were monolingual, native-English speakers and did not report any history of neurological or sleep disorders. Three participants did not complete both conditions and one participant did not complete all tasks, leaving a final sample of 32 (5 males; 27 female, age *M* = 20.94 years, *SD* = 1.53, range: 18.75 – 24.17). All participants gave informed consent. They received course credit or payment upon completion of both conditions. This study was approved by the Department of Psychology Ethics Committee at the University of York.

#### 2.1.2. Stimuli

Seventy-two stimulus pairs were devised from existing stimuli (Henderson et al., 2012; Smith et al., 2018; Weighall et al., 2017). Each pair comprised an existing familiar base word (e.g., *dolphin*) and a novel competitor (e.g., *dolpheg*). Two novel foils were also created for each pair (e.g., *dolphess* and *dolphok*) for use in the speeded recognition task. The 72 stimulus pairs were divided into four equal lists matched on written and spoken frequency, number of syllables and phonemes, as well as word length. In each condition (sleep or wake) participants were trained on one of the four lists, while another list served as a control list. Trained lists and control lists were counterbalanced across participants. All stimuli were recorded using Audacity® recording and editing software, by a female native English speaker. Stimuli lists can be found in the supplementary materials.

#### 2.1.3. Design and procedure

This experiment used a Condition (sleep vs. wake) × Session (pre-interval vs. post-interval) within-subjects design. Allocation was randomised and counterbalanced and participants completed both conditions. A schematic overview of the procedure can be found in Figure 1. Participants were asked to arrive at the laboratory at 13:00. In the sleep condition, polysomnography (PSG) was applied immediately upon arrival. In the wake condition, participants first completed a cognitive test battery (standardised scores available on the OSF). At 14:00 the pre-interval session started, in which participants in both conditions were first trained on the 18 spoken novel words via computerised training tasks. Immediately after training, participants completed three pre-tests: lexical integration was measured using a pause detection task, and explicit memory of the words was assessed using a cued recall task and a speeded recognition (old-new categorisation) task. In the sleep condition, participants were then given a 90-minute nap opportunity, while participants in the wake group watched silent movies. Silent movies were selected to reduce linguistic interference. To minimise the risk of sleep inertia influencing post-interval session performance, the post-interval session started 30 minutes after the end of the nap opportunity. This began with a Psychomotor Vigilance Test (PVT; Dinges & Powell, 1985), followed by the same three word learning post-tests: pause detection, cued recall and speeded recognition.

**Figure 1.**
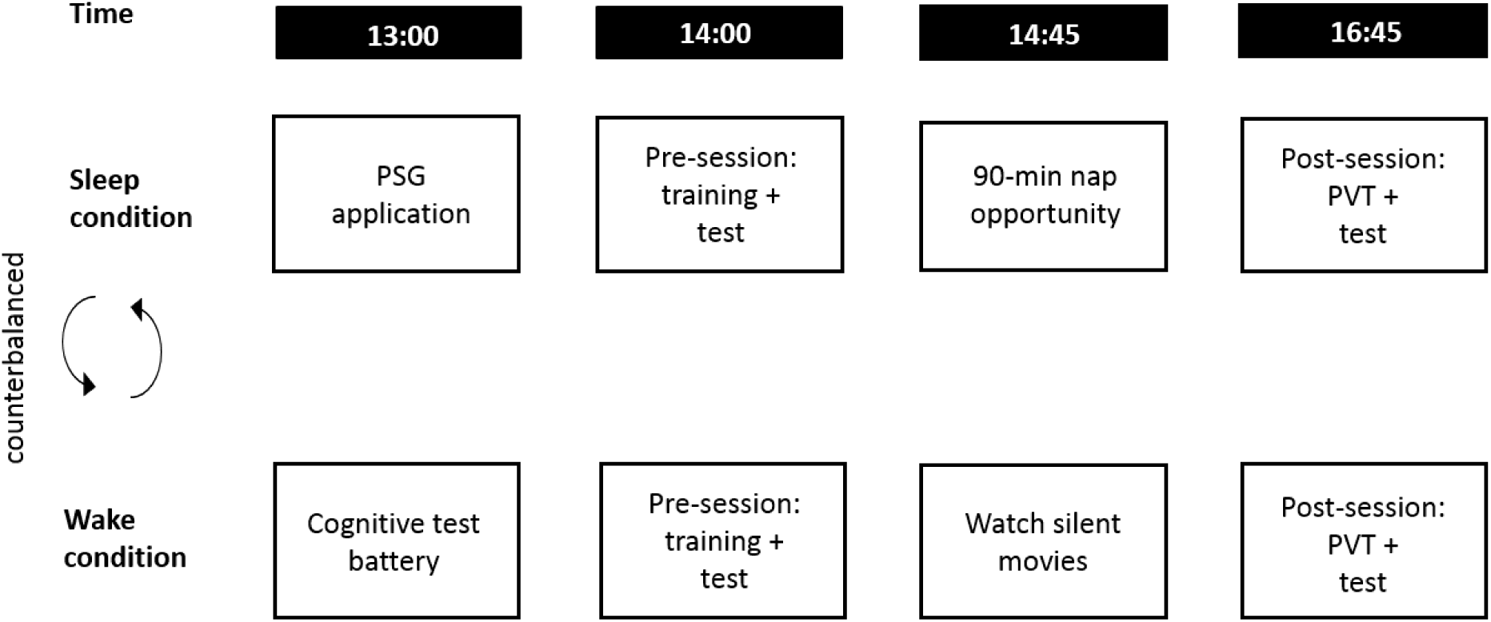
Procedure for adults. PSG = polysomnography, PVT = Psychomotor Vigilance Test.

##### 2.1.3.1. Training

Participants were exposed to 18 spoken novel words, each occurring 12 times in total across three different training tasks. In the first task, participants heard each word one by one and were instructed to repeat the word out loud. In the second task, involving phoneme monitoring, participants listened to each word and used the keyboard to indicate whether or not the word contained a pre-specified phoneme (/m/, /d/, /b/, /l/, /t/). The third task involved phoneme identification. In this task, participants again heard each word and used the mouse to select what sound each word contained from a list of four options. The three tasks alternated during the training phase, with the repetition task performed six times in total and the two phoneme tasks three times each. Stimuli were presented aurally via headphones, using Open Sesame version 3.2.4 4 (Mathôt et al., 2012). Instructions were presented both aurally and in written form (black letters on a white background, font type mono, size 18).

##### 2.1.3.2. Testing

To assess lexical integration, participants performed a *pause detection* task. In this task, participants made speeded decisions about the presence or absence of a 200 ms pause inserted into the familiar base words. A total of 72 trials were included in each test session: 18 trained words (9 with pauses), 18 untrained words (9 with pauses) and 36 fillers (18 with pauses). The words with and without pauses were counterbalanced between pre-interval session and post-interval session, and different fillers were used in each session. Ten practice trials (using a different set of familiar words) were included at the start of each session. The key dependent variables were reaction time (RT) (in ms) for correct trials, measured from the onset of the pause for both pause absent and pause present trials, and accuracy. To assess explicit memory, a cued recall and a speeded recognition task were used. In the *cued recall* task participants were presented with the first syllable of each novel word aurally (e.g., *dol-*) and were asked to recall the whole word out loud (e.g., *dolpheg*). The key dependent variable was accuracy. In the *speeded recognition* task, participants made speeded decisions about whether or not a word was learned during training. This task included 36 trials, with 18 trained words and 18 foils. The key dependent variables were RT (in ms, recorded from the beginning of the word) for correct trials and accuracy. Additionally, participants performed a 4-minute computer-based PVT (Khitrov et al., 2014) at the start of the post-interval session as an objective measure of alertness following the nap or wake period. The PVT required participants to respond with the mouse when a counter appeared in the centre of the screen, at random intervals between 2000 and 10000 ms. Number of lapses (responses with RT > 500 ms) and RT were measured. Paired-samples *t*-tests confirmed that the number of major lapses and RT did not significantly differ between conditions (number of major lapses: nap *M* = 0.03, *SD* = 0.18, wake *M* = 0.00, *SD* = 0.00, *t* = 1.00, *p* = .33; RT: nap *M* = 263.84 ms, *SD* = 28.64, wake *M* = 271.09 ms, *SD* = 24.36, *t* = −1.62, *p* = .17).

#### 2.1.4. Sleep recording and processing

Sleep was monitored in the sleep laboratory using an Embla N7000 PSG system with REMlogic software (version 3.4.0). After cleaning the scalp with NuPrep exfoliating agent (Wece and Company) gold plated electrodes were placed according to the international 10-20 system at the following locations: F3, F4, C3, C4, O1 and O2. Scalp electrodes were referenced to the contralateral mastoid (A1 or A2) and the ground electrode was placed centrally on the forehead. Electrooculography was recorded with one electrode placed next to each eye and electromyography was recorded from three electrodes, one on the chin and two under the chin. Data were sampled at 200 Hz. No filters were applied during recording.

Sleep stages were scored offline in 30-second epochs by two trained scorers according to the standard criteria of the American Academy of Sleep (AASM; Iber et al., 2007). Agreement between scores was 88.55% and any discrepancies were resolved following discussion. Time spent in each sleep stage (N1, N2, SWS and REM) was calculated. The following sleep parameters were calculated, selected based on previous research: spindle density (number of spindles per minute; fast, slow and total), sigma power (power density in the spindle frequency range (9-16 Hz); fast, slow and total) and delta power (0.3-4 Hz). Artifact rejection was performed with MATLAB, using the FieldTrip toolbox (version 10/04/2018; Oostenveld et al., 2011). Spectral analyses were conducted on artifact-free NREM epochs (high-pass filter: 0.3 Hz, low-pass filter: 30 Hz, notch filter: 50 Hz), using Fast Fourier Transformation on central channels at 12.5-16 Hz for fast sigma power and on frontal channels at 9-12.5 Hz for slow sigma power (Muehlroth et al., 2019). For delta activity (0.3-4 Hz), power was averaged across frontal and central electrodes. Irregular-resampling auto-spectral analysis was used to correct for 1/f noise (Wen & Liu, 2016). To determine NREM total spindles, a continuous wavelet transform with a morlet basis function in central and frontal channels was used at 9-16 Hz (Tsanas & Clifford, 2015). This analysis was also repeated in frequency bands 12.5-16 Hz and 9-12.5 Hz to explore both fast and slow spindles, respectively. In all cases, spindle density values reflected the mean number of spindles per minute of NREM sleep.

#### 2.1.5. Behavioural and sleep data analysis

##### 2.1.5.1. Establishing outliers for the pause detection and speeded recognition task

Accuracy rates were averaged across test sessions for the pause detection task and participants with accuracy z-scores less than −3.29 were excluded. For the speeded recognition task, a d-prime calculation based on hits (correctly identifying a learned word) and correct rejections (correctly identifying an unlearnt word) was used to determine sensitivity. Outliers were detected using z scores (+/- 3.29). For both the pause detection and speeded recognition tasks, within-subject RT outliers were classed as any trials 2.5 SDs above a participants’ mean RT. RTs < 200 ms were classed as a false response and removed. To detect outliers for individual items, accuracy rates across all test sessions were averaged separately for the pause detection and speeded recognition tasks. Items with accuracy rates 3.29 SDs below the group mean were excluded.

##### 2.1.5.2. Mixed-effects models

Data were analysed using R, with models fitted using the package *lme4* (Bates et al., 2014) and figures made using *ggplot2* (Wickham, 2016). Mixed-effects logistic regression models were used to model binary outcomes (cued recall accuracy; and if not at ceiling pause detection accuracy and speeded recognition accuracy) and linear mixed-effects models for continuous outcome data (pause detection and speeded recognition reaction times to accurate responses). For each dependent variable, fixed-effects of Condition (sleep vs. wake; +.5, −.5) and Session (pre-interval vs. post-interval; −.5, +.5) were included. For the pause detection task, Training (trained vs. not-trained items; −5 + 5) was an additional independent variable. Based on tests of normality, a transformation (log or inverse) was used to normalise the distribution of RTs if the data were found to be skewed (Brysbaert & Stevens, 2018). For the pause detection task, response latencies were measured in all cases from a marker in the digitised speech files that represented the onset of the pause in the pause-present condition (i.e. RTs measured from the same point in the speech in all cases). The fixed-effects structure was simplified using a backwards selection procedure from the maximal fixed-effects structure, using *likelihood ratio tests* (LRT) and a liberal criterion of *p* < .2 to justify inclusion. Random intercepts and slopes were justified using a liberal criterion for model improvement of *p* < .2 via LRT and added until no further model improvement could be established (Barr, 2013). The *p*-values were provided by *lmerTest* (Kuznetsova et al., 2017). After establishing the best fitting fixed-effects structure, dfbetas were calculated to identify any influential cases via the influence.ME package (Nieuwenhuis et al., 2012). Dfbetas were standardized and any participants with z-scores greater than +/-3.29 were removed from that dataset. To supplement the main analyses, following Tamminen et al. (2010), paired comparisons were run using the package *emmeans* (Lenth et al., 2018) to unpack any significant interactions and to test whether changes in performance over wake and over sleep were statistically significant. Furthermore, to determine whether lexical competition effects were present both at the pre-test and post-test, paired comparisons (trained RT vs. untrained RT) were run for each condition (wake or sleep).

##### 2.1.5.3. Correlations with sleep parameters

Correlations were run between each of the PSG sleep parameters (fast and slow spindle density, fast and slow sigma power, delta power and time in SWS) and over-nap changes in lexical competition and explicit memory. Correlations were corrected for multiple comparisons (Bonferroni corrected *p* = .002). Total sleep time (TST) was partialled out where it correlated with the sleep parameter of interest. Oscillatory power values were log transformed before correlations were performed and normality was checked for all density parameters to determine if log transformations were needed.

### 2.2. Results

#### 2.2.1. Behavioural findings

Descriptive statistics for all word learning variables can be found in Tables 1 and 2. Results tables for the mixed-effects model analyses are presented in the supplementary materials.

**Table 1.**
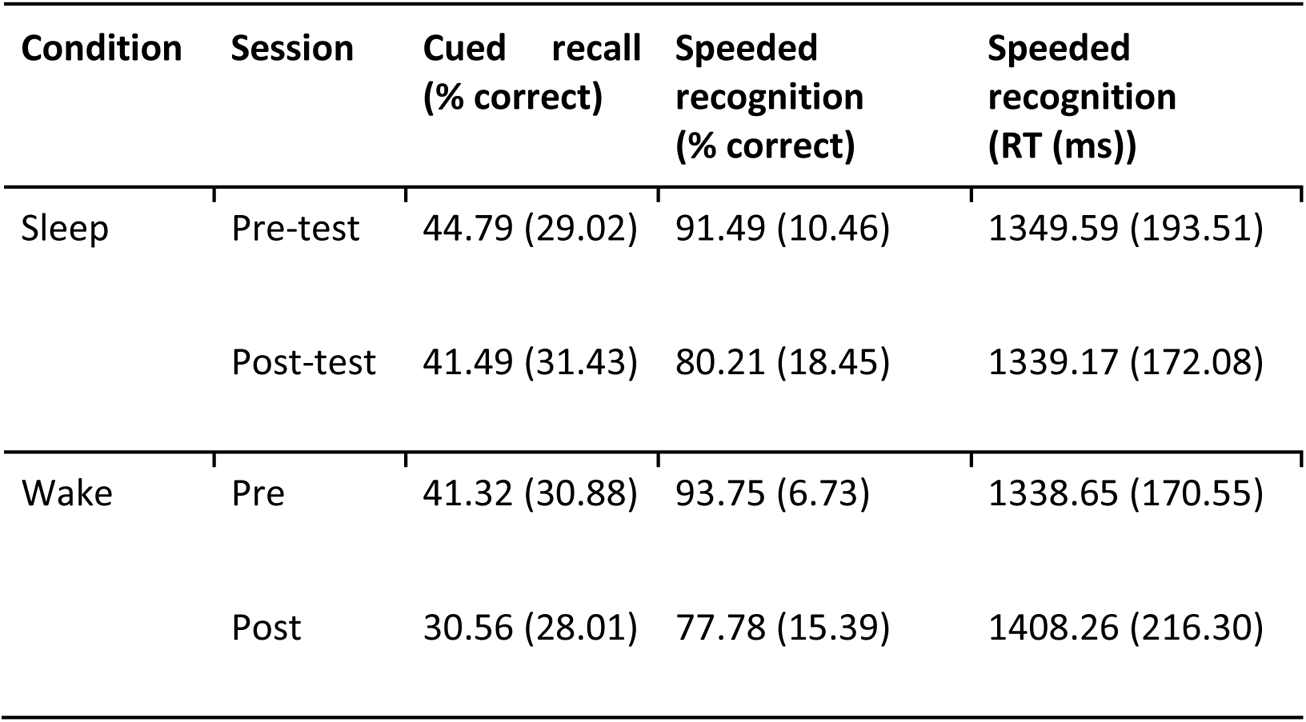
Mean (and SD) performance for the speeded recognition and cued recall tasks

| Condition | Session | Cued recall<br>(% correct) | Speeded<br>recognition<br>(% correct) | Speeded<br>recognition<br>(RT (ms)) |
| --- | --- | --- | --- | --- |
| Sleep | Pre-test | 44.79 (29.02) | 91.49 (10.46) | 1349.59 (193.51) |
|  | Post-test | 41.49 (31.43) | 80.21 (18.45) | 1339.17 (172.08) |
| Wake | Pre | 41.32 (30.88) | 93.75 (6.73) | 1338.65 (170.55) |
|  | Post | 30.56 (28.01) | 77.78 (15.39) | 1408.26 (216.30) |

##### 2.2.1.1. Cued recall

No influential cases were identified. There was a main effect of Session, such that accuracy was higher on the pre-test (*M* = 43.1%, *SD* = 29.8) than on the post-test (*M* = 36.0%, *SD* = 30; *b* = −.51, SE = .11, *p* < .001). There was also a main effect of Condition, with accuracy higher in the sleep (*M* = 43.1%, *SD* = 33.3) than in the wake condition (*M* = 35.9%, *SD* = 27.8; *b =* .64, SE = .23, *p* = .01). There was also a significant Condition × Session interaction (*b =* .57, SE = .22, *p* = .01), with performance declining significantly from the pre-test to the post-test for the wake condition (*b =* .79, SE = .16, *p* < .001), but not for the sleep condition (*b =* .22, SE = .15, *p* = .14) (Figure 2).

**Figure 2.**
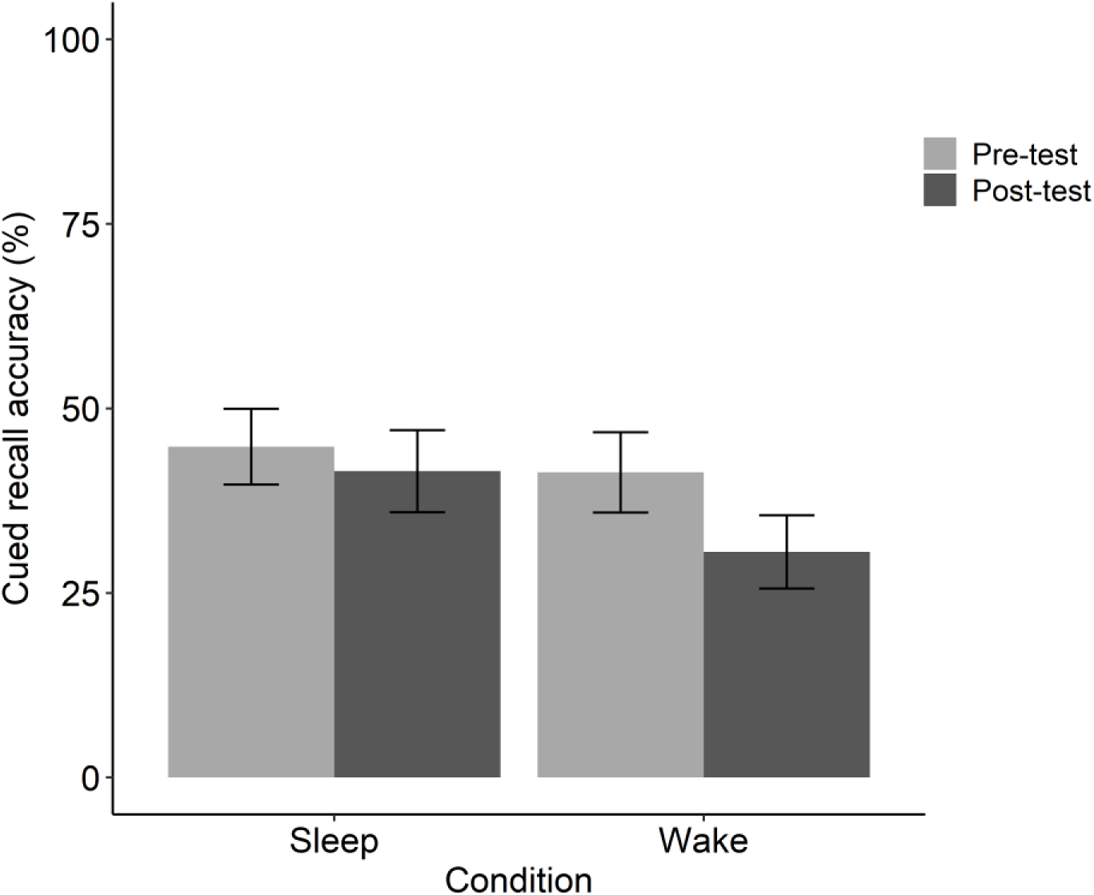
Cued recall accuracy (%) for the pre-test and post-test in the sleep and wake condition

##### 2.2.1.2. Old-new categorisation

Mixed-effects analyses were conducted on both RT and accuracy data. No outliers or influential cases were identified for either RT or accuracy. For the RT data, there was a main effect of Session, which showed that RTs were significantly slower on the post-test (*M* = 1373.72 ms, *SD* = 196.99) than on the pre-test (*M* = 1344.12 ms, *SD* = 181.02; *b =* .01, SE = .01, *p* = .01). This effect was driven by the wake group, as supported by a Condition × Session interaction (*b =* .01, SE = .01, *p* = .01). That is, there was a significant slowing of RTs from pre- to post-test for the wake condition (*b* = −.03, SE = .01, *p* < .001), but not for the sleep condition (*b* = −.00, SE = .01, *p* = .98) (see Figure 3).

**Figure 3.**
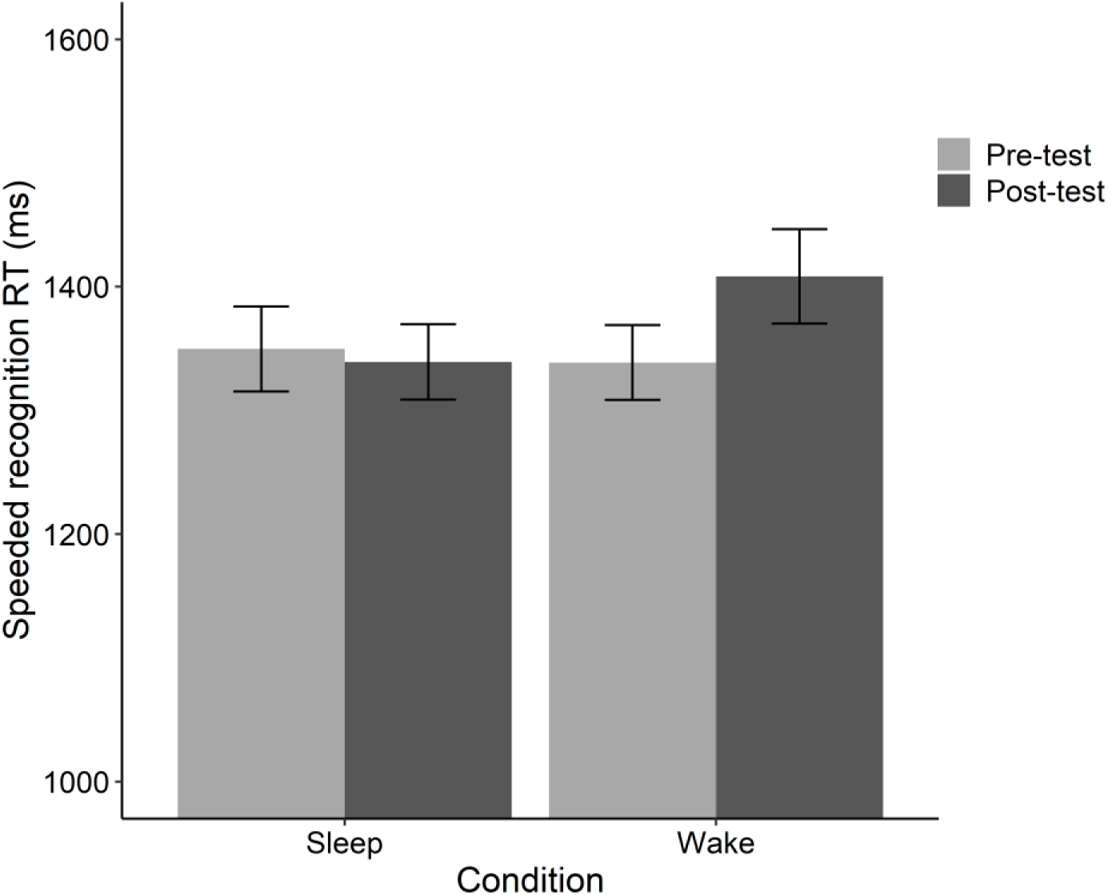
Speeded recognition reaction time (ms) for both the pre-test and post-test in the sleep and wake condition

The speeded recognition accuracy data similarly showed a main effect of Session, such that accuracy was higher on the pre-test (*M* = 92.6%, *SD* = 8.8) than on the post-test (*M* = 79.0%, *SD* = 16.9; *b =* 1.18, SE = .20, *p* < .001). As seen in Figure 4 however, the Condition × Session interaction did not reach significance (*b =* .45, SE = .28, *p* = .11).

**Figure 4.**
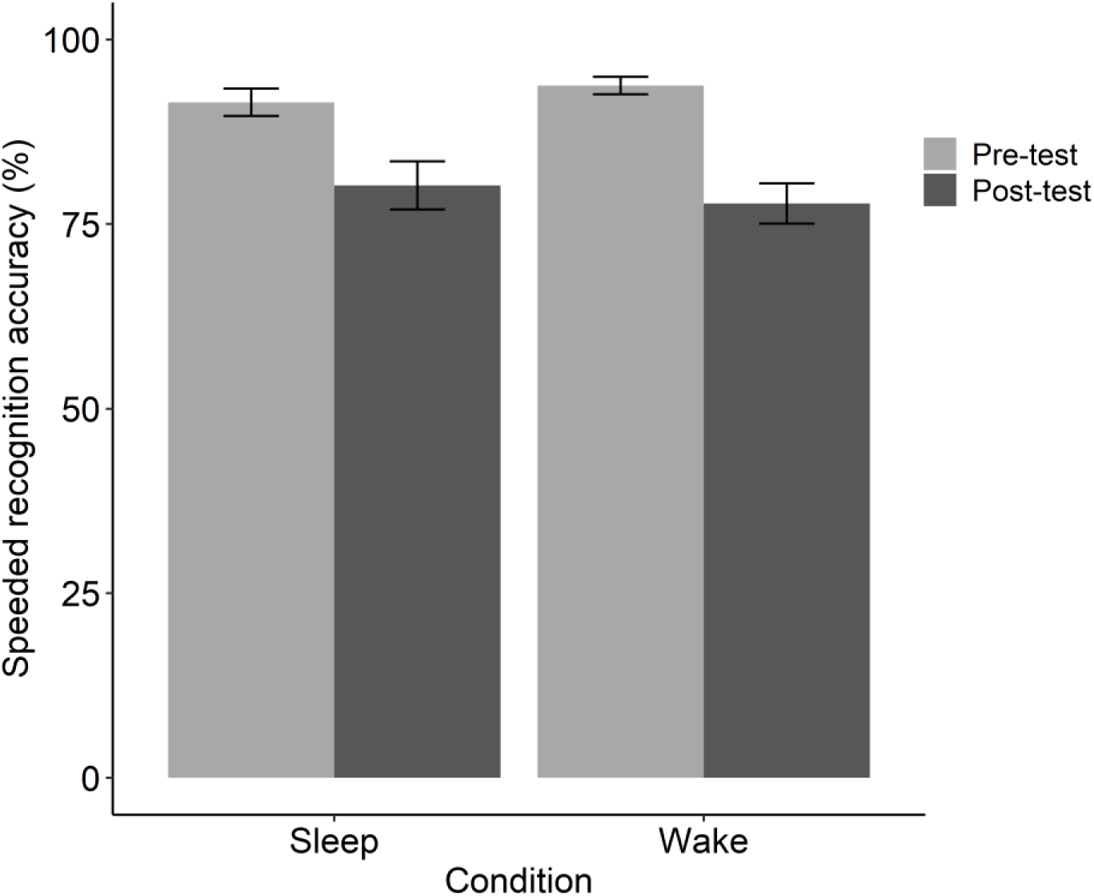
Speeded recognition accuracy (%) for the pre-test and post-test in the sleep and wake condition

##### 2.2.1.3. Pause detection

For the pause detection task, one participant was excluded due to a task administration error leaving a total sample of *N* = 31. Accuracy was at ceiling for both the sleep condition (pre-test percent accuracy: *M* = 96.77% *SD* = 4.71; post-test: *M* = 94.27%, *SD* = 6.00) and the wake condition (pre-test percent accuracy: *M* = 98.75%, *SD* = 2.76; post-test: *M* = 97.13%, *SD* = 4.27). Mixed-effects analyses were conducted on RT only. No outliers or influential cases were identified. RTs were significantly faster on the post-test (*M* = 1021.89 ms, *SD* = 139.98) than the pre-test (*M* = 978.36 ms, *SD* = 109.71; *b* = −.04, SE = .01, *p* < .001). Counter to the hypothesis, removal of the Training (i.e., trained vs. untrained) × Session and the Training × Session × Condition fixed effect did not significantly affect model fit (*p* > .2), suggesting that session had no impact on whether RTs were influenced by the words having a trained competitor or not, and crucially this was not influenced by the occurrence of a nap between sessions. RTs were significantly faster in the sleep (*M* = 1009.99 ms, *SD* = 133.33) in comparison to the wake condition (*M* = 990.26 ms, *SD* = 120.87; *b =* .02, SE = .01, *p* = .004). Counter to the hypothesis, there was no evidence of greater lexical completion in the sleep condition relative to the wake condition. Whilst the Condition × Training interaction approached significance (*b* = −.02, SE = .01, *p* = .06), exploring this with *emmeans* revealed no significant difference between trained and untrained items for the sleep condition (*b* = −.01, SE = .014, *p* = .64) and a significant difference between trained and untrained items in the wake condition (*b* = −.03, SE = .014, *p* = .035).

**Figure 5.**
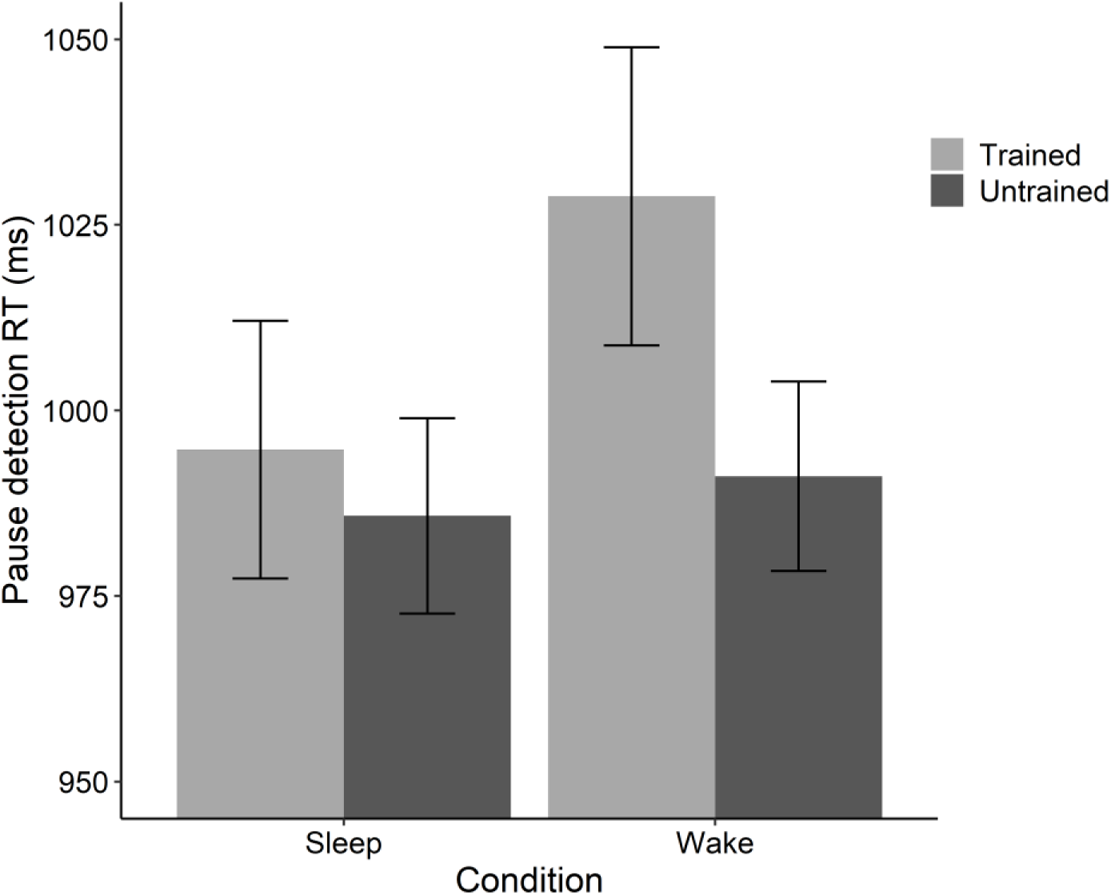
Pause detection reaction time (ms) interaction between condition (sleep vs. wake) and training (trained vs. untrained) collapsed across test sessions.

**Table 2.** Means (SD) RT and lexical competition effects (trained RT - untrained RT) for the pause detection task, for the pre-test and post-test

| Condition | Session | Trained (ms) | Untrained (ms) | Lexical competition (ms) |
| --- | --- | --- | --- | --- |
| Sleep | Pre-test | 1016.92 (142.54) | 992.79 (99.09) | 24.13 (103.72) |
|  | Post-test | 972.52 (129.24) | 978.81 (108.92) | -6.29 (100.98) |
| Wake | Pre-test | 1068.40 (184.03) | 1009.46 (114.13) | 58.94 (176.74) |
|  | Post-test | 989.32 (117.20) | 972.80 (82.79) | 16.52 (101.16) |

*Note.* N = 31

#### 2.2.2. Associations between sleep and over-nap changes in word learning

Descriptive statistics for time spent in each sleep stage can be found in Table 8. All power parameters were log transformed. All density parameters were log transformed due to normality violations. To explore whether any over-nap changes in performance (post-test - pre-test) were associated with the sleep parameters, correlations were run between over-nap change in each of the behavioural tasks and each sleep parameter: spindle density (slow and fast), sigma power (slow and fast), delta power and time in SWS. Total sleep time (TST) was partialled out for all sleep parameters except for slow spindle density. All 32 participants were included in the analyses, except for lexical competition where one participant was excluded due to missing data. As shown in Table 3, there were weak correlations between fast spindle density and over-nap changes in speeded recognition accuracy (i.e., greater improvements in accuracy associated with higher fast spindle density) and between slow spindle density and over-nap change in lexical competition (i.e., greater increase in lexical competition associated with higher slow spindle density). However, in both cases, these correlations did not survive correction for multiple comparisons (Bonferroni corrected *p* = .002).

**Table 3.** Correlations (Pearson’s r) between sleep parameters (log transformed) and over-nap change (post-test – pre-test) in cued recall accuracy, speeded recognition accuracy, speeded recognition reaction time and lexical competition in adults

|  | Over-nap<br>change in<br>cued recall<br>accuracy | Over-nap<br>change in<br>speeded<br>recognition<br>accuracy | Over-nap<br>change in<br>speeded<br>recognition<br>reaction time | Over-nap<br>change in<br>lexical<br>competition |
| --- | --- | --- | --- | --- |
| Spindle density (fast) | .159 | .379<sup>##</sup> | -.132 | -.057 |
| Spindle density (slow) | .188 | -.268 | .039 | .355<sup>#</sup> |
| Sigma power (fast) | .179 | .192 | .179 | -.074 |
| Sigma power (slow) | .135 | -.082 | -.060 | .285 |
| Delta power | -.162 | .002 | -.212 | -.012 |
| Time in SWS | -.138 | -.266 | -.099 | .144 |

Note. <sup>#</sup> $p = .049$ , <sup>##</sup> $p = .035$ (scatterplots presented in the supplementary materials), but these correlations did not survive Bonferroni correction for multiple comparisons.

### 2.3. Discussion

Experiment 1 used a within-subjects design to examine whether a day time nap (versus an equivalent period of wake) benefits novel word learning in young adults. Learning was captured via tests of explicit memory (i.e., recall and speeded recognition) and lexical integration (i.e., pause detection) that were administered before and after the nap/wake period. Sleep architecture was captured with polysomnography to examine whether over-nap changes in explicit memory or lexical integration were associated with key sleep parameters, as based on an active systems account.

Regarding explicit memory, cued recall accuracy declined significantly after a period of wake, but did not decline after a nap. Similarly, although speeded recognition accuracy was high for both groups and declined similarly over wake and sleep, speeded recognition RTs slowed down significantly over wake, but not over sleep. While this indicates a benefit of a nap over wake for novel word learning, the hypothesised enhancing effects of sleep on explicit memory (i.e., *gains* in recall and *speeding up* of recognition RTs) were not observed. Furthermore, contrary to the second hypothesis, there was no evidence for an emergence of lexical competition after the nap. Together, these data suggest that a daytime nap plays more of a protective role - guarding against forgetting - but does not actively promote the strengthening and/or lexical integration of novel word forms (consistent with the conclusions of Tamminen et al., 2017).

This interpretation aligns with the obtained pattern of correlations between key aspects of sleep known to support the consolidation of new words and over-nap changes in explicit memory and lexical integration. That is, weak correlations were found between fast and slow spindle density and over-nap changes in speeded recognition accuracy and lexical competition that did not survive correction for multiple comparisons. There was also no support for associations between sigma power, delta power or time in SWS and any of the word learning measures. Thus, counter to the third hypothesis, the over-nap changes in lexical integration and/or explicit memory were not convincingly associated with electrophysiological markers of active systems consolidation.

## 3. Experiment 2: Daytime naps and novel word learning in children

Experiment 2 examined the effect of daytime naps (versus wake) on novel word learning in children aged 10-12 years old, using the materials as Experiment 1. The pre-registered hypotheses for Experiment 2 (https://osf.io/hktwb) were largely the same as for Experiment 1, based on similar patterns of sleep-associated benefits in word learning in children (Henderson et al., 2012; James et al., 2020; Smith et al., 2018). An additional exploratory cross-experiment analysis comparing adults and children was also pre-registered. This analysis sought to examine developmental differences in (1) sleep architecture during the nap (e.g., total sleep time, time spent in each sleep stage and spindle density) and (2) over-nap changes in the word learning measures. It should be noted that while the adult study (Experiment 1) used a within-subjects design, the child study (Experiment 2) used a between-subjects design. Hence, in the cross-experiment analysis adult data from the nap condition was compared with child data from the nap group.

### 3.1. Material and methods

#### 3.1.1. Participants

Children aged 10-12 years were recruited from local primary and secondary schools, as well as through the University of York’s staff network. Due to practical and ethical considerations, separate groups of participants were recruited for the sleep (intended N = 40) and wake (intended N = 20) groups. The reason for a larger sample in the sleep group was twofold: first, to compensate for attrition due to participants not being able to fall asleep in the laboratory, and second to provide statistical power to examine associations between variability in sleep characteristics and changes in word learning across the nap. Due to COVID-19 pandemic restrictions on face-to-face contact, the final 5 participants in the wake group could not be tested. Therefore, 55 participants (24 males, age *M* = 11.79 years, *SD* = 0.89, range: 10.03 – 12.71) took part, 40 in the sleep group (age *M* = 11.92 years, *SD* = 0.91) and 15 in the wake group (age *M* = 11.55 years, *SD* = 0.80). Parental report confirmed that all participants were monolingual native English speakers and did not have any history of neurological, neurodevelopmental or sleep disorders. All parents completed the Children’s Sleep Habits Questionnaire (CSHQ; Owens et al., 2000). Independent-samples *t*-tests indicated no group differences in age and CSHQ scores (see Table 4).

**Table 4.** Group descriptives (mean (SD))

| | Sleep group | Wake group | $t$ | $p$ |
| --- | --- | --- | --- | --- |
| Age (years) | 11.90 (0.91) | 11.48 (0.78) | 1.60 | .117 |
| CSHQ (total score) | 40.90 (5.31) | 41.53 (4.87) | -0.40 | .689 |
| BPVS (standardised score) | 101.13 (11.41) | 101.53 (9.65) | -0.12 | .903 |
| BAS Matrices (standardised score) | 101.15 (16.51) | 102.47 (16.52) | -0.26 | .793 |
| Picture-vocabulary check (/ 16) | 15.43 (0.81) | 15.27 (0.88) | 0.63 | .532 |

*Note.* BPVS = British Picture Vocabulary Scale, BAS = British Ability Scales, CSHQ = Children's Sleep Habits Questionnaire

Parents were instructed to ensure their child had at least 8 hours of sleep the night before testing. On the morning of testing, they were required to get up by 7am and to not nap or consume any caffeinated products. Participants’ sleep schedules the night before the experiment were verified via verbal parental report. All participants and their parents gave written informed consent and participants were paid £60 upon completion of the study. This study was approved by the Department of Psychology Ethics Committee and the York Neuroimaging Centre Research Ethics Committee at the University of York.

#### 3.1.2. Stimuli

Thirty-two child-appropriate stimulus pairs were selected from those used in Experiment 1. The pairs were divided into two equal lists of 16 pairs, matched on age-of-acquisition (Kuperman et al., 2012), written and spoken frequency, number of syllables and phonemes, as well as word length (based on the CELEX lexical database, Kerkman et al., 1995). Participants were trained on one of the two lists, while the other list served as a control list. Trained and control lists were counterbalanced across participants. Stimuli lists can be found in the supplementary materials.

#### 3.1.3. Design and procedure

This experiment used a Group (sleep vs. wake) × Session (pre-interval vs. post-interval) between-subjects design. A schematic overview of the procedure can be found in Figure 6. Participants and their parents arrived at the laboratory at 12:30. After signing the consent forms, parents were allowed to leave. In the sleep group, PSG was then applied by one experimenter, while a second experimenter administered two standardised assessments: the British Picture Vocabulary Scale (BPVS-III; Dunn et al., 2009) the Matrices from the British Ability Scales (BAS3; Elliott & Smith, 2011). No group differences were found on these assessments (see Table 4). In the wake group, participants performed the standardised assessment upon signing the consent forms. The pre-interval session started at 13:45. Participants first filled in the Karolinska Sleepiness Scale (KSS; Åkerstedt & Gillberg, 1990) and performed a 4-minute PVT. They were then trained on the 16 spoken novel words. Immediately after training, lexical integration was measured using a pause detection task, and explicit memory of the words was assessed using a cued recall task and a speeded recognition task. Participants in the sleep group were then given a 90-minute nap opportunity, while participants in the wake group watched silent movies and played non-language based games with the experimenters (such as Bubble Shooter or Jenga). To account for sleep inertia, the post-interval session started 30 minutes after the end of the nap opportunity. The post-interval session started with the KSS and PVT, followed by the pause detection, cued recall and speeded recognition tasks. Repeated-measures ANOVAs (Group (sleep vs. wake) × Session (pre-interval vs. post-interval)) indicated no significant interactions for KSS scores, PVT number of lapses or PVT RT (all *p*s > .05). Detailed results can be found in the supplementary materials. Finally, a picture-vocabulary check was performed to ensure participants were familiar with the 16 base words. No group differences were found on the picture-vocabulary check (see Table 4).

**Figure 6.**
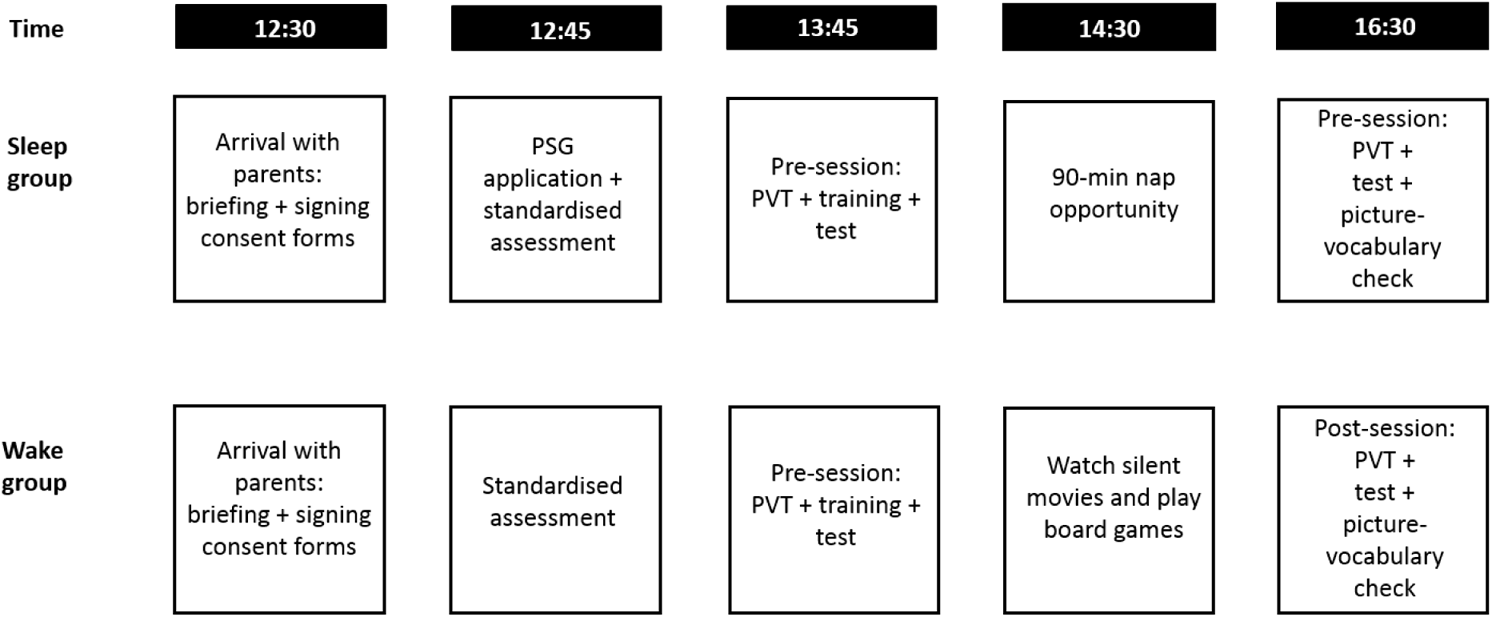
Procedure for children. PSG = polysomnography, PVT = Psychomotor Vigilance Test.

##### 3.1.3.1.Training and testing

Training and testing procedures were identical to Experiment 1, with the exception of the number of testing trials due to there being fewer trained novel words. The *pause detection* task included 64 trials in each test session: 16 trained words (8 with pauses), 16 untrained words (8 with pauses) and 32 fillers (16 with pauses). Words with and without pauses were counterbalanced between sessions, and different fillers were used in each session. Sixteen practice trials were included. The *speeded recognition* task included 32 trials, with 16 trained words and 16 foils. In the *picture-vocabulary check* at the end of the post-interval session, participants were presented with the 16 familiar base words, one by one, each accompanied by four pictures. The experimenter verbalised the word, after which the participant selected the picture that best represented the word.

#### 3.1.4. Sleep recording and processing

Sleep recording and processing were identical to Experiment 1, with the exception that two additional parietal channels (P3 and P4) were included to facilitate further exploratory analyses in preparation for future work, but were not analysed here. Agreement between scores was 95.08% and any discrepancies were resolved following discussion.

#### 3.1.5. Behavioural and sleep data analysis

Statistical analyses were identical to Experiment 1.

### 3.2. Results

Four children in the sleep group did not manage to fall asleep and were therefore excluded from all subsequent analyses, leaving a total of n = 36 in the sleep group. Descriptive statistics for all word learning variables can be found in Tables 5 and 6. Results tables for the mixed-effects model analyses are presented in the supplementary materials.

**Table 5.** Mean (SD) performance for the cued recall and speeded recognition tasks

| Group | Session | Cued recall<br>(% correct) | Speeded<br>recognition<br>(% correct) | Speeded recognition<br>(RT (ms)) |
| --- | --- | --- | --- | --- |
| Sleep | Pre-test | 25.69 (19.70) | 89.82 (13.14) | 1477.65 (198.97) |
|  | Post-test | 28.47 (23.97) | 79.82 (16.26) | 1454.46 (206.15) |
| Wake | Pre-test | 15.18 (9.72) | 87.50 (8.84) | 1482.57 (186.42) |
|  | Post-test | 9.38 (9.41) | 75.42 (10.69) | 1537.46 (185.07) |

Note. Cued recall (% correct): sleep $n = 36$ , wake $n = 14$ . Speeded recognition (% correct): sleep $n = 35$ , wake $n = 15$ . Speeded recognition (RT): sleep $n = 36$ , wake $n = 14$ .

#### 3.2.1. Behavioural findings

##### 3.2.1.1. Cued recall

One influential case was identified in the wake group, leaving n=50 in the final model (n=36 nap, n=14 wake). There was a main effect of Group, with accuracy higher in the nap group (*M* = 27.08%, *SD* = 21.83) compared to the wake group (*M* = 14.17%, *SD* = 12.17; *b* = 1.16, SE = .43, *p* = .007). In line with the predictions, there was a significant Group × Session interaction (*b =* .87, SE = .36, *p* = .017), such that there was a significant decrease in accuracy between the pre-test and post-test for the wake group (*b =* .79, SE = .33, *p* = .018), but not for the nap group (*b* = −.09, SE = .17, *p* = .617) (see Figure 7), mirroring the adult findings.

**Figure 7.**
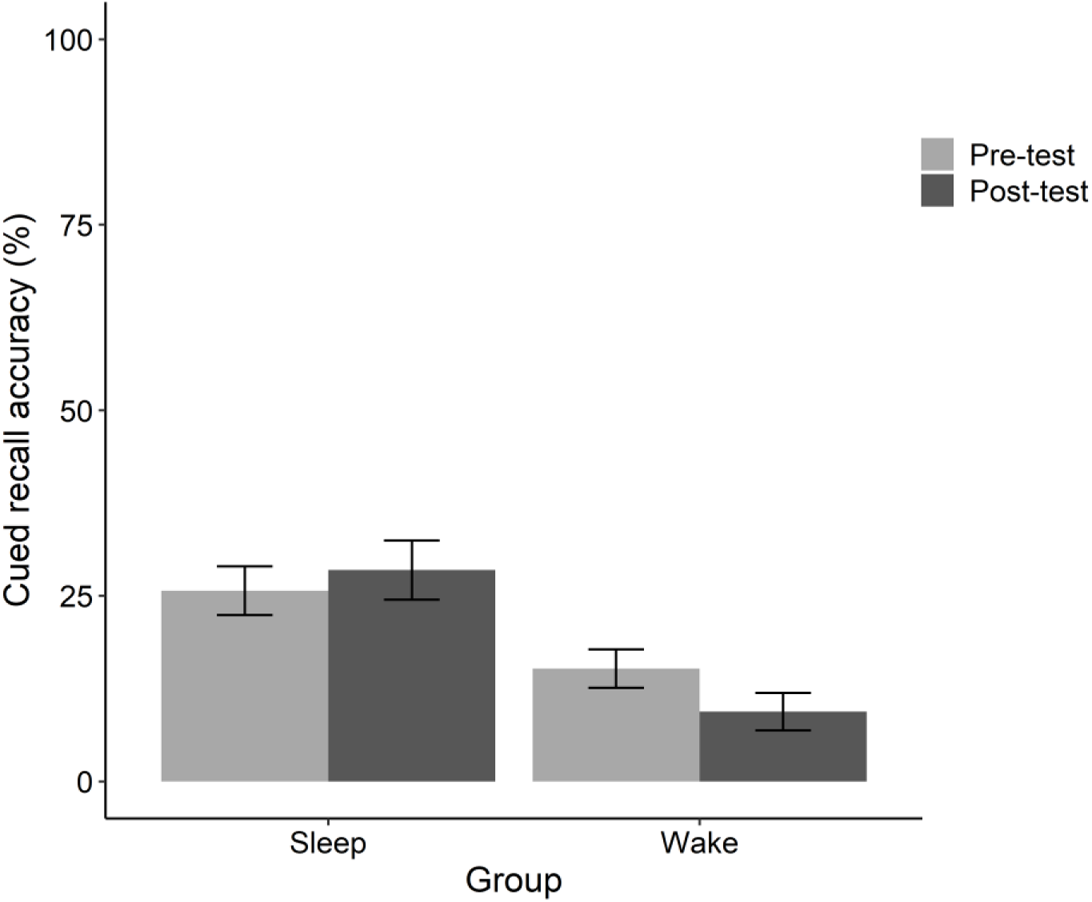
Cued recall accuracy (%) for the pre-test and post-test in the sleep and wake group

##### 3.2.1.2. Speeded recognition

Accuracy was >75% on average but not at ceiling, therefore mixed-effects analyses were conducted on both RT and accuracy data (see Table 5). For accuracy, one influential case was identified in the sleep group, leaving n=50 in the final model (n=35 nap group, n=15 wake group). There was a main effect of Session, with accuracy higher on the pre-test (*M* = 89.12%, *SD* = 11.97) than on the post-test (*M* = 78.50%, *SD* = 14.84; *b =* 0.91, SE = 0.15, *p* < .001). There was no Group × Session interaction, suggesting that a nap did not benefit post-test performance relative to an equivalent period of wake.

For speeded recognition RT, one influential case was identified in the wake group, resulting in n=50 in the final model (n=36 nap group, n=14 wake group). There were no significant main effects or interactions.

##### 3.2.1.3. Pause detection

In the wake group, one participant was detected as an outlier resulting in n=50 (n=36 nap group, n=14 wake group). Accuracy was at ceiling for both the nap group (pre-test: *M* = 96.18%, *SD* = 5.02; post-test: *M* = 96.18%, *SD* = 4.23) and the wake group (pre-test: *M* = 93.75%, *SD* = 6.00; post-test: *M* = 92.86%, *SD* = 2.80). Mixed-effects analyses were conducted on RT only. In the wake group, one further participant was detected as an influential case, resulting in n = 36 in the sleep group and n = 13 in the wake group. Children were faster on the post-test (*M* = 1124.99 ms, *SD* = 125.62) than on the pre-test (*M* = 1207.58 ms, *SD* = 149.73; *b* = −0.073, SE = 0.014, *p* < .001). The three-way interaction between Training × Session × Group was significant (*b* = 0.071, *SE* = 0.028, *p* = .013). While the interaction appeared to be driven by differences in lexical competition between the pre-test and post-test in the wake group (see Figure 8), none of the contrasts reached significance (all *p*s > .05), and the hypothesis that lexical competition would emerge after the nap was not supported.

**Figure 8.**
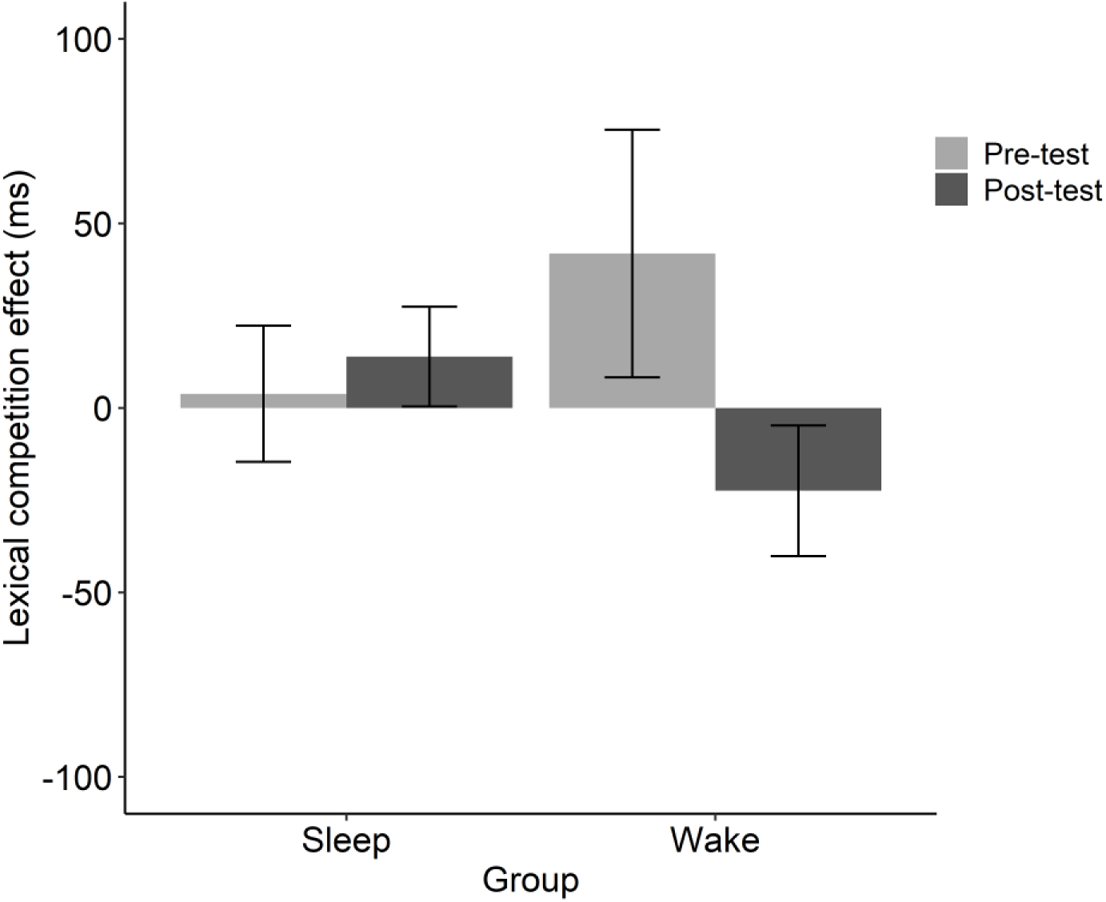
Lexical competition effect (trained - untrained reaction time) (ms) for the sleep and wake group

**Table 6.** Means (SD) RT and lexical competition effects (trained RT - untrained RT) for the pause detection task, for the pre-test and post-test

| Group | Session | Trained (ms) | Untrained (ms) | Lexical competition (ms) |
| --- | --- | --- | --- | --- |
| Sleep | Pre-test | 1211.91 (189.90) | 1208.08 (128.53) | 3.83 (110.90) |
|  | Post-test | 1141.10 (132.93) | 1127.19 (117.76) | 13.91 (81.07) |
| Wake | Pre-test | 1221.82 (110.84) | 1179.97 (120.29) | 41.85 (120.94) |
|  | Post-test | 1088.41 (122.68) | 1110.85 (134.38) | -22.44 (63.79) |

*Note.* Sleep n = 36, wake n = 13.

### 3.2.2. Associations between sleep and over-nap changes in word learning

Descriptive statistics for time spent in each sleep stage can be found in Table 8. All density parameters were log transformed due to normality violations. To explore whether any over-nap changes in performance (post-test - pre-test) were associated with the sleep parameters, correlations were run between over-nap change in each of the behavioural tasks and each sleep parameter: spindle density (slow and fast), sigma power (slow and fast), delta power and time in SWS. Total sleep time (TST) was partialled out for all sleep parameters except for slow spindle density. All participants that had slept were included in the analyses. As shown in Table 7, there was a weak correlation between slow spindle density and over-nap changes in speeded recognition reaction time (i.e., greater increase in RT associated with higher slow spindle density). However, as in Experiment 1, this correlation did not survive correction for multiple comparisons (Bonferroni corrected *p*=.002).

**Table 7.** Correlations between sleep parameters (log transformed) and over-nap change (post-test – pre-test) in cued recall accuracy, speeded recognition accuracy, speeded recognition reaction time and lexical competition in children

|  | Over-nap<br>change in<br>cued recall<br>accuracy | Over-nap<br>change in<br>speeded<br>recognition<br>accuracy | Over-nap<br>change in<br>speeded<br>recognition RT | Over-nap<br>change in<br>lexical<br>competition |
| --- | --- | --- | --- | --- |
| Spindle density (fast) | <.001 | -.030 | .119 | -.171 |
| Spindle density (slow) | -.214 | -.183 | .341<sup>#</sup> | -.153 |
| Sigma power (fast) | .158 | -.098 | .261 | -.176 |
| Sigma power (slow) | -.098 | -.277 | .327 | -.205 |
| Delta power | -.011 | .234 | -.052 | -.108 |
| Time in SWS | -.089 | .021 | -.319 | .084 |

Note. <sup>#</sup> $p = .042$ (scatterplot presented in the supplementary materials), but this correlation did not survive Bonferroni correction for multiple comparisons.

**Table 8.** Comparison of sleep parameters between children and adults.

| Sleep parameter | Adults | Children | *t* | *p* |
| --- | --- | --- | --- | --- |
|  | Mean (SD) | Mean (SD) |  |  |
| TST (minutes) | 70.20 (16.52) | 57.01 (23.03) | 2.68 | .009 |
| TST range (minutes) | 13 - 87 | 14 - 89.5 | - | - |
| N1 (%) | 10.06 (6.92) | 8.65 (7.18) | 0.82 | .414 |
| N2 (%) | 51.34 (17.63) | 39.86 (14.52) | 2.94 | .004 |
| SWS (%) | 25.65 (17.33) | 49.62 (16.91) | -5.77 | < .001 |
| REM sleep (%) | 12.96 (11.21) | 1.87 (4.34) | 5.49 | < .001 |
| Fast spindle density (raw) | 3.44 (1.75) | 1.49 (1.12) | 5.52 | < .001 |
| Slow spindle density (raw) | 8.99 (3.94) | 5.74 (3.55) | 3.58 | < .01 |
| Fast sigma power (raw) | 0.04 (0.05) | 0.01 (0.01) | 3.65 | < .01 |
| Slow sigma power (raw) | 0.04 (0.03) | 0.02 (0.03) | 1.64 | .106 |
| Delta power (raw) | 0.92 (0.10) | 0.95 (0.07) | 1.28 | .204 |

*Note.* TST = total sleep time, SWS = slow wave sleep, REM = rapid eye movement

### 3.3. Cross-experimental comparison between adults and children

#### 3.3.1. Nap characteristics

Comparisons of sleep parameters between adults and children can be found in Table 8. The adults slept significantly longer than children, on average 13.19 minutes more. Adults spent the majority of their sleep in N2, significantly more than children. Children spent most of their time asleep in SWS and their percent of SWS was significantly greater than adults. Additionally, adults spent proportionally more time in REM than children. Spindle density (fast/slow), sigma power (fast/slow) and delta power (all raw values) were compared between adults and children. Adults had significantly higher fast and slow spindle density. They also had higher fast sigma power. There were no significant differences in slow sigma power and delta power.

#### 3.3.2. Over-nap changes in word learning

Since the number of trained words was different between adults and children (18 and 16, respectively), over-nap change in % correct on the cued recall and speeded recognition task, as well as speeded recognition RT, was compared between age groups, as was over-nap change in lexical competition (see Table 9). There was a significant difference between age groups on over-nap change in cued recall: children showed a small increase in recall following the nap, while adults showed a decrease. Over-nap changes in speeded recognition or lexical competition did not differ between adults and children.

**Table 9.** Comparison of over-nap changes in word learning data between children and adults

| Variable | Adults | Children | *t* | *p* |
| --- | --- | --- | --- | --- |
|  | Mean (SD) | Mean (SD) |  |  |
| Over-nap change in cued recall accuracy (% correct) | -3.30 (10.45) | 2.78 (11.90) | -2.22 | .030 |
| Over-nap change in speeded recognition accuracy (% correct) | -11.28 (13.05) | -9.03 (10.18) | -0.80 | .427 |
| Over-nap change in speeded recognition RT (ms) | -10.42 (167.22) | -23.19 (185.39) | 0.30 | .768 |
| Over-nap change in lexical competition (ms) | -30.42 (128.29) | 10.08 (124.64) | -1.31 | .195 |

### 3.4. Discussion

#### 3.4.1. Child data

Similar to adults, there was evidence of nap-based benefits for explicit memory of new words: cued recall accuracy declined significantly after a period of wake, but not after a nap. However, counter to Experiment 1, this pattern was not mirrored in the speeded recognition data, where accuracy declined for both nap and wake groups from pre-test to post-test and RT was highly variable and produced no significant main effects or interactions. Also aligning with the adult data (but counter to the hypotheses), there was no evidence for nap-based strengthening of new word knowledge (as reflected by *maintenance* in performance rather than gains) nor was there evidence for lexical integration (owing to the absence of lexical competition at the post-test). While naps have previously been found to actively consolidate declarative memories outside of the domain of word learning in children and adults (e.g., Cairney et al., 2018; Farhadian et al., 2021; Kurdziel et al., 2013; Lokhandwala & Spencer, 2021; Tucker & Fishbein, 2008), the findings from Experiment 2, along with Experiment 1, point to a protective role of naps for explicit memory of new words, rather than naps facilitating an active systems consolidation process. Aligning with this, and similar to adults, spindle or SWS parameters were not positively associated with nap-based improvements in memory or lexical integration.

#### 3.4.2. Comparison between adults and children

Adults napped for longer than children and they spent more time in N2 and REM sleep. Spindle density (both fast and slow) and fast sigma power were also higher in adults than in children, in keeping with previous findings that developmental changes in sleep spindle characteristics are linked to the maturation of thalamo-cortical circuits (e.g., Goldstone et al., 2019; Hahn et al., 2020; Hahn et al., 2019; Zhang et al., 2021). Also in keeping with previous findings from nocturnal sleep studies (e.g., Wilhelm et al., 2013), children spent proportionally more time in SWS than adults.

Over-nap changes in speeded recognition accuracy or lexical competition did not significantly differ between children and adults. However, for cued recall accuracy, there was an interaction such that children showed an increase in recall across the nap, whereas adults showed a slight decrease. This developmental group difference in over-nap recall cannot be attributed to the greater time spent in SWS shown by the children, given the absence of correlations between over-nap change and SWS in either group. Therefore, this may suggest that there is an enhanced *protective* benefit of naps in children relative to adults, a possibility that is elaborated further in the General Discussion.

## 4. Experiment 3: Structural brain correlates of word learning in children

Experiments 1 and 2 generally lend support to theories that describe word learning as an evolving process that occurs over offline consolidation (e.g., Davis & Gaskell, 2009). However, understanding is currently lacking on the structural neural correlates of this process, particularly in childhood. As summarised earlier, DWI evidence points to three key white matter tracts that are broadly related to an array of skills related to language learning: the arcuate fasciculus (AF), left superior longitudinal fasciculus (SLF) (Romeo et al., 2018; Su et al., 2018; Urger et al., 2015), and the uncinate fasciculus (UF) (Mabbott et al., 2009; see also Lebel, Benischek, et al., 2019). The SLF and AF play a well-established role in speech and language processing, connecting the frontal cortex to the occipital, parietal and temporal cortices in the left and right hemispheres (Catani & Thiebaut de Schotten, 2008). The UF is the major white matter tract connecting frontal regions with the temporal pole, anterior temporal cortex, parahippocampal gyrus and amygdala (Von Der Heide et al., 2013), with connectivity between these regions important for declarative memory (Schmahmann et al., 2007; Squire & Zola-Morgan, 1991) and semantic processing (Holland & Lambon Ralph, 2010; Papagno et al., 2011). However, studies are yet to examine correlations between the structural integrity of these tracts and the learning and retention of new words in children.

In Experiment 3, the children who took part in Experiment 2 were invited to undergo a DWI/MRI scan, allowing us to run an exploratory analysis on the associations between fractional anisotropy (FA) in the three tracts previously broadly linked to language learning (i.e., AF, SLF, UF) and novel word learning. FA is the most commonly used measurement in DWI studies, and typically ranges from a score of 0 (indicative of isotropic movement of water molecules) to 1 (reflecting directive or anisotropic movement of water molecules, as occurs in fibre bundles such as the corpus callosum), with higher FA believed to reflect better white matter integrity. Crucially, associations with FA were examined with children’s memory of the new words (i.e., recall and recognition tasks) taken immediately after training (with children from both the wake and sleep groups combined) and also with over-nap change scores on the recall task (with children from the sleep group only) to examine neuroanatomical markers of the immediate consequences of word learning and changes over a nap interval. The recall task was selected for correlations with over-nap change, given this measure was shown at a group level to benefit from a nap (relative to wake) in Experiment 2.

### 4.1. MRI acquisition and analysis

#### 4.1.1. Participants and Scanning protocol

Participants were scanned on a separate day after the word learning procedure (nap group: range = 1 - 51 days later, M = 16.79 days later; wake group: range = 4 - 27 days later, M = 14.77 days later). Before the MRI session, participants were familiarised with the scanning procedure and environment. Participants were first shown images of the scanner, during which one of the experimenters explained the procedure. Next, they practised laying still in a mock scanner while listening to the different scanning sounds (no scan, T1 and DWI) through headphones. Diffusion-weighted images and structural data were acquired using a Siemens Magnetom Prisma 3T Scanner with a 20-channel head coil. Structural data were acquired using a T1-weighted MPRAGE whole-brain scan (TR = 2000ms, TE = 2.19ms, TI = 1040ms, flip angle = 8, FOV = 224×224mm, matrix 224×224, voxel size = 1mm^3^). Diffusion-weighted images were acquired using a spin echo planar imaging sequence with 64 diffusion encoding directions and 3 non-diffusion volumes, with the following parameters: TR = 4300ms, TE = 105ms, flip angle = 78, b = 900 s/mm^2^, FOV 240cm, matrix 160×160, 96 contiguous 1.5mm^3^ slices acquired with no gap, interleaved using a multiband acceleration factor of 4 (Moeller et al., 2010). A second diffusion scan was acquired to aid post processing for distortion correction. The same parameters were used as the 64-direction scan, but only 6 directional volumes and 1 non-diffusion volume were acquired, and the phase encode direction of acquisition was reversed. During scanning participants watched a movie of their choice projected onto a screen in the bore, with sounds delivered via MR-compatible headphones. Total scan session duration was under 30 minutes. Nine participants from Experiment 2 were not included in the analyses for the following reasons: did not get scanned (n = 3), did not fall asleep in the sleep laboratory (n = 4) or excess motion during MRI scanning (n = 2), leaving a total of n = 33 in the nap group and n = 13 in the wake group.

#### 4.1.2. Image processing

The FMRIB Software Library (FSL) data analysis suite (Smith et al., 2004) was used to perform distortion correction and to compute fractional anisotropy (FA, which is believed to measure white matter integrity), axial diffusivity (AD, reflecting axonal integrity) and radial diffusivity (RD, reflecting myelination) maps. TOPUP was used to estimate distortion in the diffusion images using the 64-direction diffusion dataset and the reverse phase encoded dataset. Once computed, the warp field map correction was applied to the 64-direction dataset for further analysis. Nonlinear registration was performed on all FA images in order to align them into the 1×1×1mm FMRIB58_FA standard space. Next, the nonlinear transformed images were aligned into the 1×1×1mm MNI152 standard space. The mean FA and the FA skeleton were then generated from the mean of the subjects. The mean FA skeleton was thresholded at FA > 0.2 to select major fibre bundles only. FA, AD and RD maps were created based on data from all 46 participants, as well as for the sleep group separately.

#### 4.1.3. Regions of interest

To examine the relationship between white matter integrity and language learning, the following white matter tracts were selected as regions of interest (ROIs): arcuate fasciculus (AF), superior longitudinal fasciculus (SLF) and uncinate fasciculus (UF). Left and right hemispheres were analysed separately.

#### 4.1.4. Tract-based spatial statistics (TBSS)

FSL’s randomize tool (using 5000 permutations) (Smith et al., 2006) was used to carry out voxelwise statistical analyses of FA, AD and RD data in above specified ROIs. Results were corrected for multiple comparisons using threshold-free cluster enhancement (TFCE; Smith & Nichols, 2009), which then allowed for estimation of cluster sizes corrected for family-wise error (FWE; *p* < .05)

To identify if the ROIs were involved in language learning in the current sample, FA, AD and RD measures in all participants were correlated with the following variables from the pre-interval test session of the word learning task: cued recall accuracy score, speeded recognition accuracy score and speeded recognition reaction time. Additionally, change scores (where positive scores reflect an increase in recall from pre-test to post-test, while negative scores reflect forgetting following the nap) were examined in the sleep group separately for variables from the word learning task in which a significant difference was found between the sleep and wake group, in this case cued recall. For the significant correlations (as indicated by the TBSS analysis) the FA, AD or RD values were extracted for the significant voxels (*p* < .05) and averaged for each participant. Next, Pearson’s correlations were run between each DWI index (FA, AD or RD) and the relevant corresponding word learning variable. Only the FA analyses, which was the primary measure of interest, are reported here. Analyses of AD and RD, as well as analyses with sleep variables, can be found on the OSF (https://osf.io/wpd8v/).

### 4.2 Results

#### 4.2.1. Relationship between FA and word learning measures on the pre-interval test session

Correlations were run between RA in the ROIs and word learning measures at the pre-interval test session, to index white-matter correlates of the immediate consequences of word learning. No significant correlations were found between pre-test cued recall scores and FA in any of the ROIs. Pre-test speeded recognition accuracy scores and RT correlated positively with FA in the bilateral AF (accuracy left AF: *r* = 0.414, *p* = .004, accuracy right AF: *r* = 0.440, *p* = .002; RT left AF: *r* = 0.514, *p* < .001; RT right AF: *r* = 0.417, *p* = .004) and the right UF (accuracy: *r* = 0.528, *p* < .001, RT: *r* = 0.528, *p* < .001) (see Figures 9 and 10).

**Figure 9.**
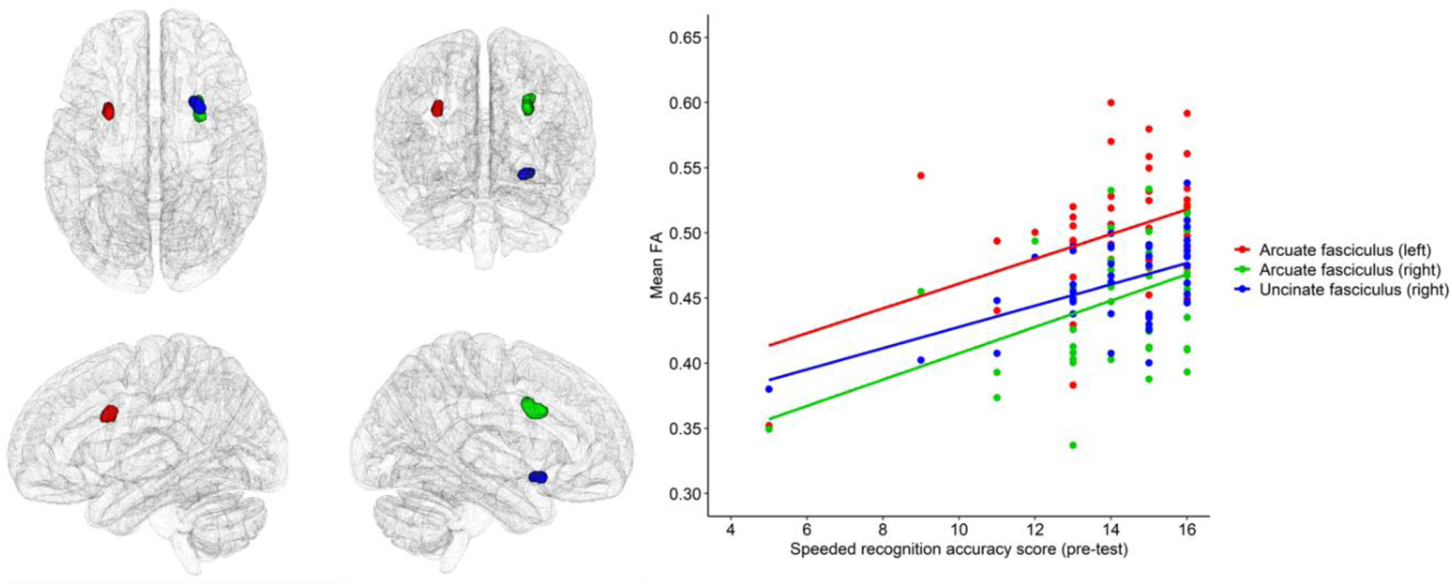
Correlation between FA and speeded recognition accuracy score on the pre-test in the bilateral arcuate fasciculus and the right uncinate fasciculus

**Figure 10.**
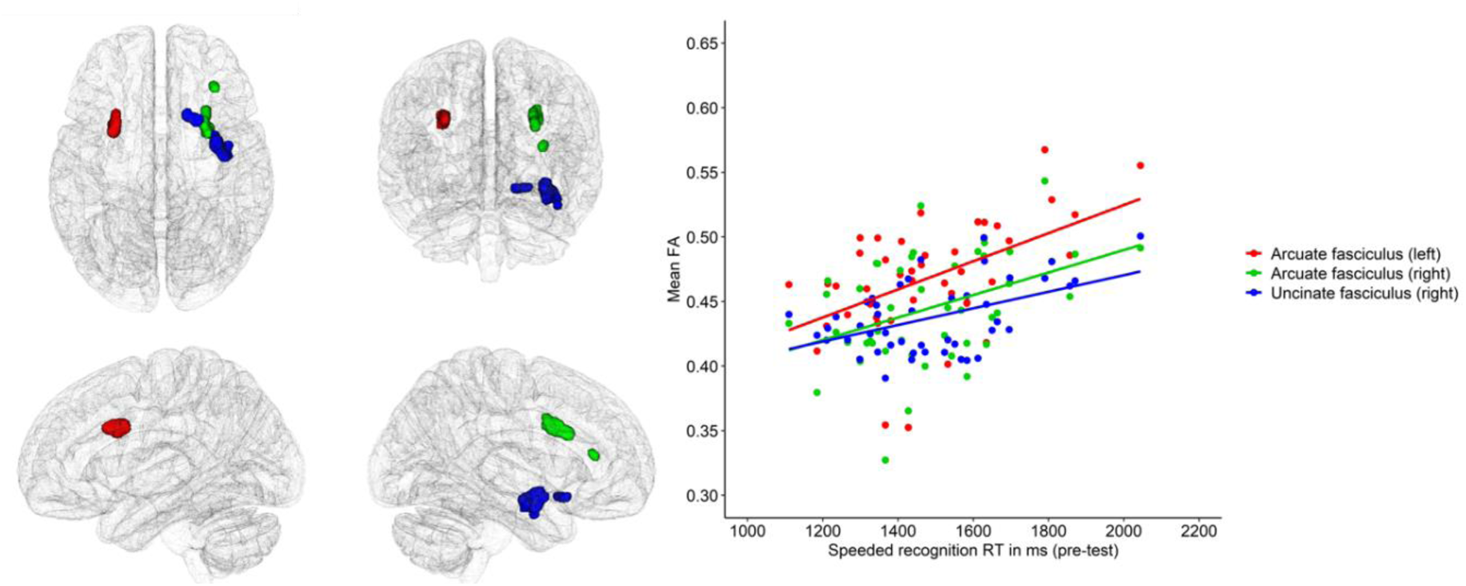
Correlation between FA and speeded recognition reaction time on the pre-test in the bilateral arcuate fasciculus and the right uncinate fasciculus in all participants

#### 4.2.2. Relationship between FA and over-nap change in cued recall

As reported in section 3.2.1.1, there was a significant Group × Session interaction for cued recall accuracy, driven by a decrease in accuracy between the pre-test and post-test in the wake group, but not the nap group. Therefore, change scores in cued recall accuracy (an index of off-line changes in novel word representations) were correlated with the FA measures in the nap group (n = 33). Particularly given the exploratory nature of the analyses, correlations were not run for the wake group to the relatively small number of participants in that group (n = 13). Cued recall accuracy change scores correlated positively with FA in the right AF, such that participants with larger increase in recall accuracy following the nap had higher FA in the right AF (see Figure 12).

**Figure 12.**
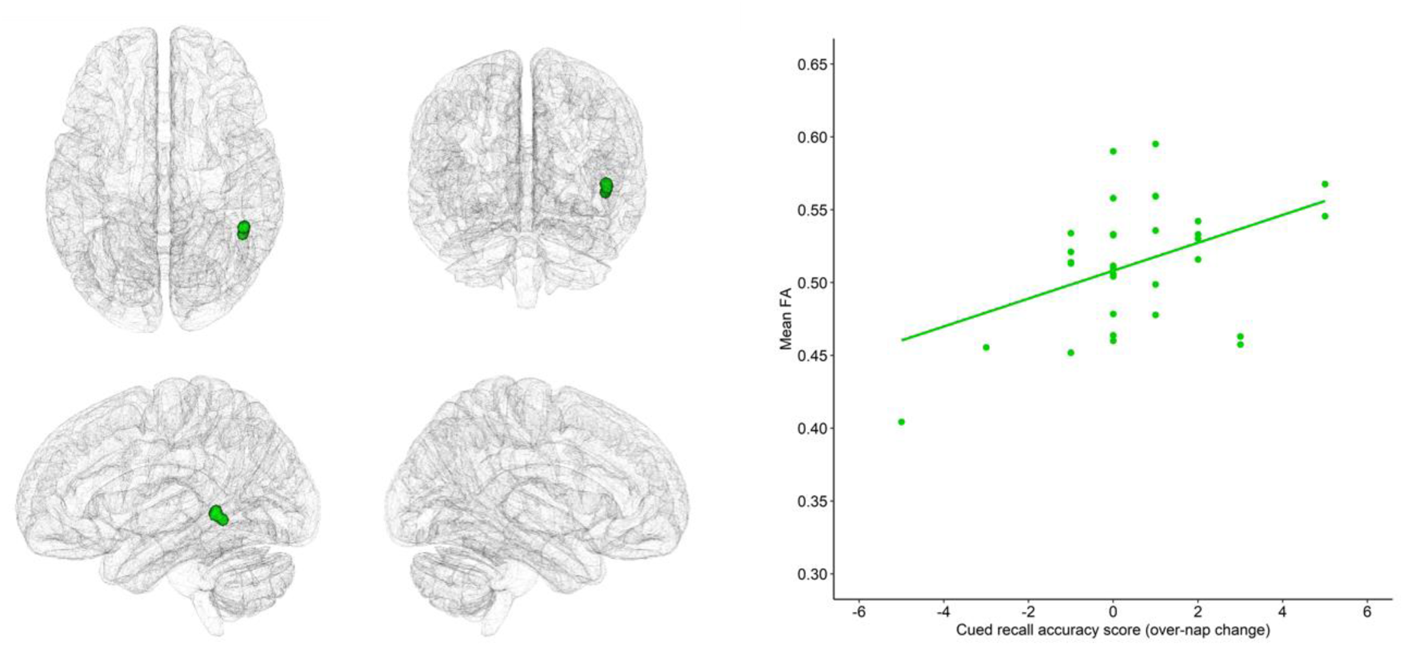
Correlation between FA in the right arcuate fasciculus and over-nap change in cued recall accuracy in the sleep group

### 4.3. Discussion

Experiment 3 comprised an exploratory analysis of white matter correlates of word learning. When considering the immediate consequences of word learning, speeded recognition accuracy scores significantly correlated with FA in the bilateral AF and the right UF, such that children with higher speeded recognition accuracy had higher FA in these white matter tracts. These correlations are broadly in line with previous studies linking these tracts to various language measures (e.g., Catani & Thiebaut de Schotten, 2008; Mabbott et al., 2009; Romeo et al., 2018; Su et al., 2018; Urger et al., 2015; Von Der Heide et al., 2013). It should be noted that these correlations were coupled with positive correlations between speeded recognition response time and FA in the same tracts, most likely due to a speed-accuracy trade-off (i.e., children who responded more accurately also responded more slowly).

Despite observing no correlations between pre-test cued recall accuracy and FA in any of the ROIs, over-nap changes in cued recall accuracy were significantly (and positively) associated with FA in the right AF. Children with higher FA in this tract had greater increases in explicit memory of the novel words following the nap. Given that a comparable analysis of the wake group was not feasible, the behavioural changes cannot be exclusively attributed to sleep. Nevertheless, it can be concluded that *offline changes* in memory for novel words are associated with white matter integrity of the right AF.

## 5. General Discussion

This study examined whether daytime naps, like periods of nocturnal sleep, benefit the consolidation of novel words, as well as their integration into the lexical network. This was investigated in two age groups: young adults and children aged 10-12 years old. To examine if children’s word learning benefits more from naps than adults (similar to previously reported enhanced effects of nocturnal sleep in children e.g., see James et al., 2017 for a review), the two age groups were compared on nap sleep parameters and on change in word learning performance across the nap. Finally, an exploratory experiment conducted for the child group aimed to identify white matter correlates of word learning.

### 5.1. Daytime naps play a protective role in novel word learning

For children and adults, a 90-minute daytime nap led to the maintenance of explicit memory of novel words, as measured via a cued recall task, whereas an equivalent wake period led to forgetting. Whilst adults were also slower at recognising novel words following wake compared to sleep, there was no effect of sleep on speeded recognition accuracy for either age group (potentially due to higher levels of initial performance leaving less room for improvement). Furthermore, neither age group demonstrated evidence of lexical integration (as indexed by lexical competition between the known and trained words) following a nap. This falls in contrast to studies examining overnight sleep and word learning in children and adults, which demonstrated sleep-associated improvements in explicit memory, as well as engagement in competition (Fletcher et al., 2020; Smith et al., 2018; Tamminen et al., 2013; Tamminen et al., 2010; Weighall et al., 2017). There were also no statistically significant correlations between over-nap changes in explicit memory or lexical integration and sleep parameters known to support systems consolidation (Born & Wilhelm, 2012; Diekelmann & Born, 2010; Rasch & Born, 2013). Notably, previous studies of both children and adults, using similar novel word learning paradigms, have found improvements in recall of novel words following a full night of sleep, evidence of lexical integration, and associations between these changes and sleep spindle parameters (Henderson et al., 2012; Smith et al., 2018; Tamminen et al., 2010). Thus, while the current results support a protective effect of daytime naps relative to wake for explicit memory in children and adults, there was no evidence that naps play an active role in strengthening and integrating newly learned words. Possibly then, a 90-minute nap is beneficial for maintaining new memories of novel words, perhaps through guarding against forgetting that might occur during wake as a consequence of interference (Wixted, 2004), but not for supporting active systems consolidation, at least at the age groups of focus and when daytime naps are not habitual (consistent with Tamminen et al., 2017). This account explains why retention of information after a nap was still superior to wake and why the nap did not promote strengthening or integration of the new words.

One possibility is that sleep parameters need to reach a particular threshold before active systems consolidation is observed. Consistent with this, Piosczyk et al. (2013) found that a daytime nap promoted memory consolidation of word-pairs in 16-year-old females, but only when the nap was characterised by high sigma power. Another explanation for why effects that emerge across nocturnal sleep were not seen here, is because levels of REM sleep were low. Indeed, it has been shown that vocabulary learning might benefit from REM sleep that occurs after SWS during an afternoon nap (Batterink et al., 2017). During nocturnal sleep, SWS and REM sleep alternate cyclically, with SWS naturally followed by REM sleep (Rasch & Born, 2013). Various theoretical models have proposed that SWS and REM play complementary roles in memory consolidation. Active systems accounts argue that whilst systems consolidation occurs during SWS (repeatedly activating newly encoded memories and driving the integration of new memories into the network of pre-existing long-term memories), REM sleep acts to stabilize the transformed memories by enabling undisturbed synaptic consolidation (Diekelmann & Born, 2010). According to the sequential hypothesis (e.g., Ambrosini & Giuditta, 2001), SWS works to eliminate non-adaptive memory traces, with subsequent REM sleep strengthening the remaining traces. Walker and Stickgold (2010) further postulate that REM sleep plays an integrative function, forming rich associative links between new and existing knowledge. Thus, nocturnal sleep, rich with SWS and REM sleep, might therefore lead to more opportunities for spontaneous reactivation of newly learned words. For example, a recent study demonstrated that toddlers that had both a daytime nap and night time sleep benefitted the most for generalisation of novel categories, compared to those that did not nap or were tested 4-hrs after the nap (Werchan et al., 2021).

This raises an important point: Post-training naps may be particularly powerful for enhancing longer-term retention when they are combined with nocturnal sleep. It has been shown that when 6-7-year-olds napped soon after mirror discrimination training this led to significantly after gains in reading fluency than compared to not napping after training, even when performance was tested after nocturnal sleep (Torres et al., 2021). These results support the claim that the delay between training and sleep onset is a critical factor, particularly in younger learners (e.g., Backhaus et al., 2008; James et al., 2020; Walker et al., 2020), and crucially, that the protective effects of a nap may work in combination with nocturnal sleep, as a useful memory enhancer.

### 5.2. Developmental differences in nap architecture and over-nap change in word learning

Despite adults napping for longer than children and spending more time in N2 and REM sleep, children exhibited significantly (and proportionally) more SWS. This is consistent with previous reports of greater nocturnal SWS in children in this age range compared to adults (e.g., Wilhelm et al., 2013). Spindle density (both fast and slow) and sigma power (fast only) were also higher in adults than in children. Higher sigma power in adults aligns with previous findings demonstrating that sigma frequency increases linearly with age (Zhang et al., 2021) and plateaus in adulthood (Purcell et al., 2017). Spindle density, on the other hand, increases from childhood to adolescence, peaks at age 15 (Zhang et al., 2021), and then declines from middle to late adulthood (Purcell et al., 2017). Given that the children tested in the current study were 10-12 years old, it is likely that they had not yet reached peak spindle density, which could explain the higher spindle density found in the young adults. Overall then, the comparison between adult and child nap architecture is in line with previous studies of nocturnal sleep (Wilhelm et al., 2013; Zhang et al., 2021).

Behaviourally, children showed an increase in cued recall accuracy across the nap, while this decreased slightly in adults. This difference in over-nap change in cued recall between the two age groups was significant, albeit small numerically. Whilst children spent more time in SWS than adults, the group difference in recall cannot be simply attributed to increased opportunities for spontaneous reactivation of the new memories during SWS in children, given that there were no correlations between over-nap change in cued recall and time in SWS or delta power in either group. It is possible that the children’s levels of alertness benefited more from the nap than adults, accounting for their better post-nap performance for cued recall. However, as children were slower on the post-nap PVT than adults (but had similar major lapses)^2^, we think this to be an unlikely explanation.

The enhanced protective role of naps for children relative to adults (coupled with no correlations with SWS or spindle parameters) raises the possibility that previously reported enhanced benefits of consolidation over nocturnal sleep in children might not be solely due to extra amounts of SWS. It has been claimed that reactivation of newly acquired memories can occur without sleep, during periods of wakeful rest, and memory benefits associated with wakeful rest have been attributed to the absence of interfering information (Dewar et al., 2012; Dewar et al., 2014; Mednick et al., 2011; Wixted, 2004). However, whilst studies have demonstrated benefits of wakeful rest for word-pair recall in children (e.g., Martini et al., 2021), no studies to our knowledge have directly addressed whether children show greater effects of wakeful rest relative to adults. It also remains unclear whether the present findings could be explained by developmental differences in susceptibility to proactive or retroactive interference. Preliminary findings suggest that retroactive interference effects remain robust across development but that adults can be more resilient to the effects of retroactive interference than children (e.g., Darby & Sloutsky, 2015; Fatania & Mercer, 2017), however, the literature is sparse on whether there are developmental differences in proactive or retroactive interference in the context of novel word learning. For example, if children are at increased risk of interference when attempting to recall newly learned information, one possibility is that a nap allows the developing system to be “reset”, decreasing susceptibility to such interference and enhancing performance. However, this remains highly speculative. Aside from having pedagogical value, addressing these questions will be theoretically important in deepening our understanding of the various routes by which newly learned words become established in long-term language networks.

### 5.3. Structural brain correlates of word learning in children

The final contribution of this study was to explore the structural neuroanatomical correlates of the word learning process, both immediately after learning and short-term changes that occurred over a nap opportunity. Faster speeded recognition of novel words immediately after learning was associated with higher fractional anisotropy in the bilateral AF and the right UF. This suggests that children with higher white matter integrity in these brain regions are also better at recognising new words immediately after training. These results align with previous research on various aspects of language use and language learning (Farah et al., 2020; Lebel, Benischek, et al., 2019; Mabbott et al., 2009; Romeo et al., 2018; Su et al., 2018), solidifying the importance of these tracts as part of the developing language learning network. Previous studies have linked the integrity of these tracts to a broad range of language measures in children (e.g., expressive language ability, Farah et al., 2020; auditory-verbal memory, Mabbott et al., 2009; conversational experience, Romeo et al., 2018; rate of vocabulary growth, Su et al., 2018); however, the present results importantly add to this existing literature by demonstrating a role for the AF and UF in recognising novel words, pointing to an underlying learning mechanism for the speeded recognition of new word forms.

Another finding of note here was although cued recall accuracy at immediate testing was not related to any of the ROIs, over-nap change in recall accuracy was positively associated with FA in the right AF. This finding supports the role of the right AF in the retention of novel words, as opposed to initial encoding processes. The fact that these correlations emerged in the right hemisphere is perhaps surprising in the context of the commonly reported left lateralization of the language network (e.g., Romeo et al., 2018; Sreedharan et al., 2015; Su et al., 2018; Urger et al., 2015). However, there have been other studies demonstrating associations with language learning in the right hemisphere (e.g., Farah et al., 2020; Romeo et al., 2018; Sreedharan et al., 2015; Su et al., 2018). Takashima et al. (2019) found that when processing newly learned words, young children (8-10 years old) had greater right lateralized activation in the inferior frontal gyrus (IFG), while older children (14-16 years old) had greater activation in the left IFG. This is in line with studies demonstrating a shift from right to left lateralization as a result of cognitive maturation, as well as increased lexical knowledge or familiarity (Sugiura et al., 2011; Szaflarski et al., 2006; Takashima et al., 2019). Possibly, as children get older and their lexical knowledge increases, a similar shift to left lateralised structural correlates of word learning would be observed.

In sum, models of word learning distinguish between processes that help us to quickly acquire new words from the environment and processes that support their long-term consolidation into existing vocabulary, suggesting that variability might emerge at multiple stages of new word acquisition (see James et al., 2017). Here, these models are expanded by identifying structural neuroanatomical correlates of the word learning process, both immediately after learning and over a nap, importantly identifying different neuroanatomical correlates at each point. Such correlates will have use for future studies which track the time course of word learning over longer time periods, including over cumulative periods of sleep. In adults, white matter integrity has been shown to be associated with offline gains in motor memory, with nocturnal sleep spindle density mediating this relationship (Mander et al., 2017; Vien et al., 2019). Therefore, the identification of white matter correlates of both the immediate consequences of word learning and changes over a nap in children informs future examinations of whether these white matter correlates dictate the functional influence of sleep spindles on the consolidation of new words.

### 5.4. Limitations and conclusions

There are a number of limitations that need to be considered when interpreting the present results. Although the sample sizes used here were comparable to previous studies reporting associations between word learning and sleep in children and adults (e.g., Smith et al., 2018; Tamminen et al., 2010) the sample sizes were nevertheless small, which may have contributed to the weak correlations. Notably, the sample size was particularly small (n = 15) in the children’s wake group (owing to data collection ceasing during the COVID-19 pandemic), and whilst this does not influence the cross-development differences or the correlations between over-nap changes in performance and white matter integrity (which were based on the nap groups only; n = 32 adults and n = 36 children), it will be important to replicate the sleep-wake differences in the children. Furthermore, the adult and child studies utilised within- and between-subjects designs respectively, limiting the extent to which the datasets can be directly compared.

To conclude, daytime naps maintained explicit memory of novel words in children and adults, supporting a protective effect of naps on memory. Unlike previously reported benefits of nocturnal sleep, there was no evidence that naps benefitted lexical integration or initiated active systems consolidation of new word representations. Of course, naps may be able to promote active systems consolidation under different circumstances (e.g., when naps are longer). Consistent with previously observed age-related differences in nocturnal sleep, children had more SWS during the nap than adults, while adults had higher spindle density and fast sigma power. Children showed a slight over-nap increase in cued recall whereas recall declined over the nap slightly for adults, but these changes did not correlate with any sleep parameters. Therefore, rather than this enhanced behavioural effect in children being driven by sleep, children may instead benefit more from an offline period that is free from further incoming information than adults. Finally, this study provides preliminary insight into the white matter structures that support the immediate and longer-term consequences of word learning in children, identifying structural integrity of the bilateral AF and the right UF as two key tracts, and opening up future lines of research that delve deeper into the role of these tracts in support the word learning process. Together, these findings demonstrate how models of word learning and sleep-associated memory consolidation can be advanced via the lenses of developmental and neuroimaging data.

## Acknowledgements

The authors wish to thank Rebecca Anderson, Gabrielle Appleby, Amanda Olsson, Mohreet Rauni, Anna Rodgerson and Francesca Wilson, as well as the team at York Neuroimaging Centre, for their assistance with data collection. This work was supported by the Economic and Social Research Council, grant number ES/N009924/1.

## OSF links

Study 1: https://osf.io/ehw5z/?view_only=b471cfc06d884bcb8fd63d1a9a27ed4f

Adults pre-registration: https://osf.io/63p2e

Study 2: https://osf.io/wpd8v/?view_only=21cf82dd7608474d908b5b9d5b1c2786

Children pre-registration: https://osf.io/hktwb

## Footnotes

1 Sigma power was featured in the pre-registration analysis plan, but was omitted (in error) from the hypotheses.

2 RT: t(66) = −8.25, p = .009; major lapses: t(66) = −1.26, p = .214)

## Notes

https://osf.io/ehw5z/?view_only=b471cfc06d884bcb8fd63d1a9a27ed4f

https://osf.io/63p2e

https://osf.io/wpd8v/?view_only=21cf82dd7608474d908b5b9d5b1c2786

https://osf.io/hktwb

### Competing Interest Statement

The authors have declared no competing interest.

## References

Åkerstedt, T., & Gillberg, M. (1990). Subjective and objective sleepiness in the active individual. International Journal of Neuroscience, 52, 29–37. doi:10.3109/00207459008994241

Ambrosini, M. V., & Giuditta, A. (2001). Learning and sleep: The sequential hypothesis. Sleep Medicine Reviews, 5(6), 477–490. doi:10.1053/smrv.2001.0180

Armstrong, R., Arnott, W., Copland, D. A., McMahon, K., Khan, A., Najman, J. M., & Scott, J. G. (2017). Change in receptive vocabulary from childhood to adulthood: associated mental health, education and employment outcomes. International Journal of Language & Communication Disorders, 52(5), 561–572. doi:10.1111/1460-6984.12301

Bakker-Marshall, I., Takashima, A., Schoffelen, J. M., van Hell, J. G., Janzen, G., & McQueen, J. M. (2018). Theta-band oscillations in the middle temporal gyrus reflect novel word consolidation. Journal of Cognitive Neuroscience, 30(5), 621–633. doi:10.1162/jocn_a_01240

Barakat, M., Doyon, J., Debas, K., Vandewalle, G., Morin, A., Poirier, G., … Carrier, J. (2011). Fast and slow spindle involvement in the consolidation of a new motor sequence. Behavioural Brain Research, 217(1), 117–121. doi:10.1016/j.bbr.2010.10.019

Barr, D. J. (2013). Random effects structure for testing interactions in linear mixed-effects models. Frontiers in Psychology, 4, 328. doi:10.3389/fpsyg.2013.00328

Bates, D., Maechler, M., Bolker, B., & Walker, S. (2014). lme4: Linear mixed-effects models using ‘Eigen’ and S4. R Package, Version 1(7).

Batterink, L. J., Westerberg, C. E., & Paller, K. A. (2017). Vocabulary learning benefits from REM after slow-wave sleep. Neurobiology of Learning and Mememory, 144, 102–113. doi:10.1016/j.nlm.2017.07.001

Bishop, D.V., Barry, J.G., & Hardiman, M.J. (2012). Delayed retention of new word-forms is better in children than adults regardless of language ability: A factorial two-way study. PLoS ONE, 7, e37326.

Born, J., & Wilhelm, I. (2012). System consolidation of memory during sleep. Psychological Research, 76(2), 192–203. doi:10.1007/s00426-011-0335-6

Breitenstein, C., Jansen, A., Deppe, M., Foerster, A. F., Sommer, J., Wolbers, T., & Knecht, S. (2005). Hippocampus activity differentiates good from poor learners of a novel lexicon. Neuroimage, 25(3), 958–968. doi:10.1016/j.neuroimage.2004.12.019

Brown, H., Weighall, A., Henderson, L. M., & Gaskell, M. G. (2012). Enhanced recognition and recall of new words in 7- and 12-year-olds following a period of offline consolidation. Journal of Experimental Child Psychology, 112(1), 56–72. doi:10.1016/j.jecp.2011.11.010

Brysbaert, M., & Stevens, M. (2018). Power analysis and effect size in mixed effects models: A tutorial. Journal of Cognition, 1(1), 9. doi:10.5334/joc.10

Cabral, T., Mota, N. B., Fraga, L., Copelli, M., McDaniel, M. A., & Ribeiro, S. (2018). Post-class naps boost declarative learning in a naturalistic school setting. npj Science of Learning, 3, 14. doi:10.1038/s41539-018-0031-z

Cairney, S. A., Guttesen, A. A. V., El Marj, N., & Staresina, B. P. (2018). Memory consolidation is linked to spindle-mediated information processing during sleep. Current Biology, 28(6), 948–954 e944. doi:10.1016/j.cub.2018.01.087

Campbell, I. G., & Feinberg, I. (2009). Longitudinal trajectories of non-rapid eye movement delta and theta EEG as indicators of adolescent brain maturation. Proceedings of the National Academy of Sciences of the United States of America, 106(13), 5177–5180. doi:10.1073/pnas.0812947106

Casey, B. J., Tottenham, N., Liston, C., & Durston, S. (2005). Imaging the developing brain: What have we learned about cognitive development? Trends in Cognitive Sciences, 9(3), 104–110. doi:10.1016/j.tics.2005.01.011

Catani, M., & Thiebaut de Schotten, M. (2008). A diffusion tensor imaging tractography atlas for virtual in vivo dissections. Cortex, 44(8), 1105–1132. doi:10.1016/j.cortex.2008.05.004

Chatburn, A., Coussens, S., Lushington, K., Kennedy, D., Baumert, M., & Kohler, M. (2013). Sleep spindle activity and cognitive performance in healthy children. SLEEP, 36(2), 237–243. doi:10.5665/sleep.2380

Darby K., Sloutsky V. M. Proactive and retroactive interference effects in development. In: Knauff M., Pauen M., Sebanz N., Wachsmuth I., editors. Proceedings of the 35th Annual Conference of the Cognitive Science Society. Austin, TX: Cognitive Science Society; 2013. pp. 2130–2135.

Davis, M. H., Di Betta, A. M., Macdonald, M. J., & Gaskell, M. G. (2009). Learning and consolidation of novel spoken words. Journal of Cognitive Neuroscience, 21(4), 803–820. doi:10.1162/jocn.2009.21059

Davis, M. H., & Gaskell, M. G. (2009). A complementary systems account of word learning: Neural and behavioural evidence. Philosophical Transansactions of the Royal Society, 364(1536), 3773–3800. doi:10.1098/rstb.2009.0111

De Gennaro, L., & Ferrara, M. (2003). Sleep spindles: An overview. Sleep Medicine Reviews, 7(5), 423–440. doi:10.1016/S1087-0792(02)00116-8

Dewar, M., Alber, J., Butler, C., Cowan, N., & Della Sala, S. (2012). Brief wakeful resting boosts new memories over the long term. Psychological Science, 23(9), 955–960. doi:10.1177/0956797612441220

Dewar, M., Alber, J., Cowan, N., & Della Sala, S. (2014). Boosting long-term memory via wakeful rest: Intentional rehearsal is not necessary, consolidation is sufficient. PLoS One, 9(10), e109542. doi:10.1371/journal.pone.0109542

Diekelmann, S., Biggel, S., Rasch, B., & Born, J. (2012). Offline consolidation of memory varies with time in slow wave sleep and can be accelerated by cuing memory reactivations. Neurobiology of Learning and Memory, 98(2), 103–111. doi:10.1016/j.nlm.2012.07.002

Diekelmann, S., & Born, J. (2010). The memory function of sleep. Nature Review Neuroscience, 11(2), 114–126. doi:10.1038/nrn2762

Dinges, D. F., & Powell, J. W. (1985). Microcomputer analyses of performance on a portable, simple visual RT task during sustained operations. Behavior, Research Methods, Instruments, & Computers, 17(6), 652–655.

Dumay, N., & Gaskell, M. G. (2007). Sleep-associated changes in the mental representation of spoken words. Psychological Science, 18, 35–39. doi:10.1111/j.1467-9280.2007.01845.x

Dunn, D. M., Dunn, L. M., Styles, B., & Sewell, J. (2009). The British Picture Vocabulary Scale III - 3rd Edition. London: GL Assessment.

Elliott, C. D., & Smith, P. (2011). The British Ability Scales, 3rd Edition. London: GL Assessment.

Farah, R., Tzafrir, H., & Horowitz-Kraus, T. (2020). Association between diffusivity measures and language and cognitive-control abilities from early toddler’s age to childhood. Brain Structure & Function, 225(3), 1103–1122. doi:10.1007/s00429-020-02062-1

Farhadian, N., Khazaie, H., Nami, M., & Khazaie, S. (2021). The role of daytime napping in declarative memory performance: a systematic review. Sleep Medicine, 84 134–141. doi:10.1016/j.sleep.2021.05.019

Fletcher, F. E., Knowland, V., Walker, S., Gaskell, M. G., Norbury, C., & Henderson, L. M. (2020). Atypicalities in sleep and semantic consolidation in autism. Developmental Science, 23(3), e12906. doi:10.1111/desc.12906

Gaskell, M. G., & Dumay, N. (2003). Lexical competition and the acquisition of novel words. Cognition, 89(2), 105–132. doi:10.1016/s0010-0277(03)00070-2

Giganti, F., Arzilli, C., Conte, F., Toselli, M., Viggiano, M. P., & Ficca, G. (2014). The effect of a daytime nap on priming and recognition tasks in preschool children. SLEEP, 37(6), 1087–1093. doi:10.5665/sleep.3766

Goldstone, A., Willoughby, A. R., de Zambotti, M., Clark, D. B., Sullivan, E. V., Hasler, B. P., … Baker, F. C. (2019). Sleep spindle characteristics in adolescents. Clinical Neurophysiology, 130(6), 893–902. doi:10.1016/j.clinph.2019.02.019

Hahn, M. A., Heib, D., Schabus, M., Hoedlmoser, K., & Helfrich, R. F. (2020). Slow oscillation-spindle coupling predicts enhanced memory formation from childhood to adolescence. Elife, 9, 1–21. doi:10.7554/eLife.53730

Hahn, M. A., Joechner, A. K., Roell, J., Schabus, M., Heib, D. P., Gruber, G., … Hoedlmoser, J. (2019). Developmental changes of sleep spindles and their impact on sleep-dependent memory consolidation and general cognitive abilities: A longitudinal approach. Developmental Science, 22(1), e12706. doi:10.1111/desc.12706

Henderson, L., Powell, A., Gaskell, M. G., & Norbury, C. (2014). Learning and consolidation of new spoken words in autism spectrum disorder. Developmental Science, 17(6), 858–871. doi:10.1111/desc.12169

Henderson, L. M., Weighall, A. R., Brown, H., & Gaskell, M. G. (2012). Consolidation of vocabulary is associated with sleep in children. Developmental Science, 15(5), 674–687. doi:10.1111/j.1467-7687.2012.01172.x

Hofstetter, S., Friedmann, N., & Assaf, Y. (2017). Rapid language-related plasticity: microstructural changes in the cortex after a short session of new word learning. Brain Structure & Function, 222(3), 1231–1241. doi:10.1007/s00429-016-1273-2

Holland, R., & Lambon Ralph, M. A. (2010). The anterior temporal lobe semantic hub is a part of the language neural network: selective disruption of irregular past tense verbs by rTMS. Cerebral Cortex, 20(12), 2771–2775. doi:10.1093/cercor/bhq020

Houston, J., Allendorfer, J., Nenert, R., Goodman, A. M., & Szaflarski, J. P. (2019). White matter language pathways and language performance in healthy adults across ages. Frontiers in Neuroscience, 13, 1185. doi:10.3389/fnins.2019.01185

Iber, C., Ancoli-Israel, S., Chesson, A. L. J., & Quan, S. F. (2007). The AASM manual for the scoring of sleep and associated events: rules, terminology, and technical specification (1 ed.). Westchester, IL: American Academy of Sleep Medicine.

James, E., Gaskell, M. G., & Henderson, L. M. (2019). Offline consolidation supersedes prior knowledge benefits in children’s (but not adults’) word learning. Developmental Science, 22(3), e12776. doi:10.1111/desc.12776

James, E., Gaskell, M. G., & Henderson, L. M. (2020). Sleep-dependent consolidation in children with comprehension and vocabulary weaknesses: it’ll be alright on the night? Journal of Child Psychology and Psychiatry. doi:10.1111/jcpp.13253

James, E., Gaskell, M. G., Weighall, A., & Henderson, L. (2017). Consolidation of vocabulary during sleep: The rich get richer? Neuroscience and Biobehavioral Reviews, 77, 1–13. doi:10.1016/j.neubiorev.2017.01.054

Ji, X., Li, J., & Liu, J. (2019). The relationship between midday napping and neurocognitive function in early adolescents. Behavioral Sleep Medicine, 17(5), 537–551. doi:10.1080/15402002.2018.1425868

Kapnoula, E. C., Packard, S., Gupta, P., & McMurray, B. (2015). Immediate lexical integration of novel word forms. Cognition, 134, 85–99. doi:10.1016/j.cognition.2014.09.007

Kerkman, H., Piepenbrock, R., Baayen, R. H., & van Rijn, H. (1995). The CELEX lexical database. Philadelphia, PA: Linguistic Data Consortium.

Khitrov, M. Y., Laxminarayan, S., Thorsley, D., Ramakrishnan, S., Rajaraman, S., Wesensten, N. J., & Reifman, J. (2014). PC-PVT: a platform for psychomotor vigilance task testing, analysis, and prediction. Behavior Research Methods, 46(1), 140–147. doi:10.3758/s13428-013-0339-9

Khundrakpam, B. S., Reid, A., Brauer, J., Carbonell, F., Lewis, J., Ameis, S., … Brain Development Cooperative, G. (2013). Developmental changes in organization of structural brain networks. Cereb Cortex, 23(9), 2072–2085. doi:10.1093/cercor/bhs187

Kuperman, V., Stadthagen-Gonzalez, H., & Brysbaert, M. (2012). Age-of-acquisition ratings for 30,000 English words. Behavior Research Methods, 44(4), 978–990. doi:10.3758/s13428-012-0210-4

Kurdziel, L., Duclos, K., & Spencer, R. M. (2013). Sleep spindles in midday naps enhance learning in preschool children. Proceedings of the National Academy of Sciences of the United States of America, 110(43), 17267–17272. doi:10.1073/pnas.1306418110

Kurdziel, L., Mantua, J., & Spencer, R. M. C. (2017). Novel word learning in older adults: A role for sleep? Brain & Language, 167, 106–113. doi:10.1016/j.bandl.2016.05.010

Kurth, S., Jenni, O. G., Riedner, B. A., Tononi, G., Carskadon, M. A., & Huber, R. (2010). Characteristics of sleep slow waves in children and adolescents. SLEEP, 33(4), 475–480.

Kuznetsova, A., Brockhoff, P. B., & Christensen, R. H. B. (2017). lmerTest Package: Tests in minear mixed effects Models. Journal of Statistical Software, 82(13). doi:10.18637/jss.v082.i13

Landi, N., Malins, J. G., Frost, S. J., Magnuson, J. S., Molfese, P., Ryherd, K., … Pugh, K. R. (2018). Neural representations for newly learned words are modulated by overnight consolidation, reading skill, and age. Neuropsychologia, 111, 133–144. doi:10.1016/j.neuropsychologia.2018.01.011

Lau, E. Y. Y., McAteer, S., Leung, C. N. W., Tucker, M. A., & Li, C. (2018). Beneficial effects of a daytime nap on verbal memory in adolescents. Journal of Adolescence, 67, 77–84. doi:10.1016/j.adolescence.2018.06.004

Lebel, C., Benischek, A., Geeraert, B., Holahan, J., Shaywitz, S., Bakhshi, K., & Shaywitz, B. (2019). Developmental trajectories of white matter structure in children with and without reading impairments. Developmental Cognitive Neuroscience, 36, 100633. doi:10.1016/j.dcn.2019.100633

Lebel, C., Treit, S., & Beaulieu, C. (2019). A review of diffusion MRI of typical white matter development from early childhood to young adulthood. NMR Biomed, 32(4), e3778. doi:10.1002/nbm.3778

Lemos, N., Weissheimer, J., & Ribeiro, S. (2014). Naps in school can enhance the duration of declarative memories learned by adolescents. Frontiers in Systems Neuroscience, 8, 103. doi:10.3389/fnsys.2014.00103

Lenth, R., Singmann, H., Love, J., Buerkner, P., & Herve, M. (2018). emmeans: Estimated Marginal Means, aka Least-Squares Means (Version R Package, Version 1.3.2). Retrieved from https://CRAN.R-project.org/package=emmeans

Lindsay, S., & Gaskell, M. G. (2013). Lexical integration of novel words without sleep. Journal of Experimental Psychology: Learning, Memory, and Cognition, 39(2), 608–622. doi:10.1037/a0029243

Lokhandwala, S., & Spencer, R. M. C. (2021). Slow wave sleep in naps supports episodic memories in early childhood. Developmental Science, 24(2), e13035. doi:10.1111/desc.13035

Mabbott, D. J., Rovet, J., Noseworthy, M. D., Smith, M. L., & Rockel, C. (2009). The relations between white matter and declarative memory in older children and adolescents. Brain Research, 1294, 80–90. doi:10.1016/j.brainres.2009.07.046

Mander, B. A., Zhu, A. H., Lindquist, J. R., Villeneuve, S., Rao, V., Lu, B., … Walker, M. P. (2017). White matter structure in older adults moderates the benefit of sleep spindles on motor memory consolidation. Journal of Neuroscience, 37(48), 11675–11687. doi:10.1523/JNEUROSCI.3033-16.2017

Marshall, L., & Born, J. (2007). The contribution of sleep to hippocampus-dependent memory consolidation. Trends in Cognitive Sciences, 11(10), 442–450. doi:10.1016/j.tics.2007.09.001

Martini, M., Martini, C., & Sachse, P. (2021). Brief period of post-encoding wakeful rest supports verbal memory retention in children aged 10–13 years. Current Psychology, 40(5), 2341–2348. doi:10.1007/s12144-019-0156-0

Mathôt, S., Schreij, D., & Theeuwes, J. (2012). OpenSesame: An open-source, graphical experiment builder for the social sciences. Behavior Research Methods, 44(2), 314–324. doi:10.3758/s13428-011-0168-7

Mattys, S. L., & Clark, J. H. (2002). Lexical activity in speech processing: evidence from pause detection. Journal of Memory and Language, 47 343–359.

McClelland, J. L., McNaughton, B. L., & O’Reilly, R. C. (1995). Why there are complementary learning-systems in the hippocampus and neocortex: Insights from the successes and failures of connectionist models of learning and memory. Psychological Review, 102, 419–457. doi:10.1037/0033-295X.102.3.419

McMurray, B., Munson, C., & Tomblin, J. B. (2014). Individual differences in language ability are related to variation in word recognition, not speech perception: Evidence from eye movements. Journal of Speech, Language, and Hearing Research, 57(4), 1344–1362. doi:10.1044/2014_JSLHR-L-13-0196

McMurray, B., Samelson, V. M., Lee, S. H., & Tomblin, J. B. (2010). Individual differences in online spoken word recognition: Implications for SLI. Cognitive Psychology, 60(1), 1–39. doi:10.1016/j.cogpsych.2009.06.003

Mednick, S. C., Cai, D. J., Shuman, T., Anagnostaras, S., & Wixted, J. T. (2011). An opportunistic theory of cellular and systems consolidation. Trends in Neurosciences, 34(10), 504–514. doi:10.1016/j.tins.2011.06.003

Miller, D. J., Duka, T., Stimpson, C. D., Schapiro, S. J., Baze, W. B., McArthur, M. J., … Sherwood, C. C. (2012). Prolonged myelination in human neocortical evolution. Proceedings of the National Academy of Sciences of the United States of America 109(41), 16480–16485. doi:10.1073/pnas.1117943109

Moeller, S., Yacoub, E., Olman, C. A., Auerbach, E., Strupp, J., Harel, N., & Ugurbil, K. (2010). Multiband multislice GE-EPI at 7 tesla, with 16-fold acceleration using partial parallel imaging with application to high spatial and temporal whole-brain fMRI. Magnetic Resonance in Medicine, 63(5), 1144–1153. doi:10.1002/mrm.22361

Mölle, M., Bergmann, T. O., Marshall, L., & Born, J. (2011). Fast and slow spindles during the sleep slow oscillation: disparate coalescence and engagement in memory processing. SLEEP 34(10), 1411–1421.

Mölle, M., Marshall, L., Gais, S., & Born, J. (2004). Learning increases human electroencephalographic coherence during subsequent slow sleep oscillations. Proceedings of the National Academy of Sciences of the United States of America, 101(38), 13963–13968.

Muehlroth, B. E., Sander, M. C., Fandakova, Y., Grandy, T. H., Rasch, B., Shing, Y. L., & Werkle-Bergner, M. (2019). Precise slow oscillation-spindle coupling promotes memory consolidation in younger and older adults. Scientific Reports, 9(1), 1940. doi:10.1038/s41598-018-36557-z

Nation, K. (2014). Lexical learning and lexical processing in children with developmental language impairments. Philosophical Transactions of the Royal Society, 369(1634), 20120387. doi:10.1098/rstb.2012.0387

Nieuwenhuis, R., Pelzer, B., & te Grotenhuis, M. (2012). influence.ME: Tools for detecting influential data in mixed effects models (R package version 0.9). Enschede: R. Nieuwenhuis. Retrieved from http://CRAN.R-project.org/package=influence.ME

Ohayon, M. M., Carskadon, M. A., Guilleminault, C., & Vitiello, M. V. (2004). Meta-analysis of quantitative sleep parameters from childhood to old age in healthy individuals: developing normative dleep values across the human lifespan. SLEEP, 27(7), 1255–1273.

Oostenveld, R., Fries, P., Maris, E., & Schoffelen, J. M. (2011). FieldTrip: Open source software for advanced analysis of MEG, EEG, and invasive electrophysiological data. Computational Intelligence and Neuroscience, 2011, 156869. doi:10.1155/2011/156869

Owens, J. A., Spirito, A., & McGuinn, M. (2000). The Children’s Sleep Habits Questionnaire (CSHQ): Psychometric properties of a survey instrument for school-aged children. SLEEP, 23(8), 1043–1051.

Papagno, C., Miracapillo, C., Casarotti, A., Romero Lauro, L. J., Castellano, A., Falini, A., … Bello, L. (2011). What is the role of the uncinate fasciculus? Surgical removal and proper name retrieval. Brain, 134(Pt 2), 405–414. doi:10.1093/brain/awq283

Peiffer, A., Brichet, M., De Tiege, X., Peigneux, P., & Urbain, C. (2020). The power of children’s sleep - Improved declarative memory consolidation in children compared with adults. Scientific Reports, 10(1), 9979. doi:10.1038/s41598-020-66880-3

Piosczyk, H., Holz, J., Feige, B., Spiegelhalder, K., Weber, F., Landmann, N., … Nissen, C. (2013). The effect of sleep-specific brain activity versus reduced stimulus interference on declarative memory consolidation. Jounral of Sleep Research, 22(4), 406–413. doi:10.1111/jsr.12033

Plihal, W., & Born, J. (1997). Effects of early and late nocturnal sleep on declarative and procedural memory. Journal of Cognitive Neuroscience, 94, 534–547.

Purcell, S. M., Manoach, D. S., Demanuele, C., Cade, B. E., Mariani, S., Cox, R., … Stickgold, R. (2017). Characterizing sleep spindles in 11,630 individuals from the National Sleep Research Resource. Nature Communications, 8, 15930. doi:10.1038/ncomms15930

Rasch, B., & Born, J. (2013). About sleep’s role in memory. Physiological Reviews, 93(2), 681–766. doi:10.1152/physrev.00032.2012

Romeo, R. R., Segaran, J., Leonard, J. A., Robinson, S. T., West, M. R., Mackey, A. P., … Gabrieli, J. D. E. (2018). Language Exposure Relates to Structural Neural Connectivity in Childhood. Journal of Neuroscience, 38(36), 7870–7877. doi:10.1523/JNEUROSCI.0484-18.2018

Schmahmann, J. D., Pandya, D. N., Wang, R., Dai, G., D’Arceuil, H. E., de Crespigny, A. J., & Wedeen, V. J. (2007). Association fibre pathways of the brain: Parallel observations from diffusion spectrum imaging and autoradiography. Brain, 130(Pt 3), 630–653. doi:10.1093/brain/awl359

Schmithorst, V. J., Wilke, M., Dardzinski, B. J., & Holland, S. K. (2005). Cognitive functions correlate with white matter architecture in a normal pediatric population: a diffusion tensor MRI study. Human Brain Mapping, 26(2), 139–147. doi:10.1002/hbm.20149

Shtyrov, Y., Nikulin, V. V., & Pulvermuller, F. (2010). Rapid cortical plasticity underlying novel word learning. Journal of Neuroscience, 30(50), 16864–16867. doi:10.1523/JNEUROSCI.1376-10.2010

Smalle, E. H. M., Page, M. P. A., Duyck, W., Edwards, M., & Szmalec, A. (2018). Children retain implicitly learned phonological sequences better than adults: a longitudinal study. Developmental Science, 21(5), e12634.

Smith, F. R. H., Gaskell, M. G., Weighall, A. R., Warmington, M., Reid, A. M., & Henderson, L. M. (2018). Consolidation of vocabulary is associated with sleep in typically developing children, but not in children with dyslexia. Developmental Science, 21(5), e12639. doi:10.1111/desc.12639

Smith, S. M., Jenkinson, M., Johansen-Berg, H., Rueckert, D., Nichols, T. E., Mackay, C. E., … Behrens, T. E. (2006). Tract-based spatial statistics: voxelwise analysis of multi-subject diffusion data. Neuroimage, 31(4), 1487–1505. doi:10.1016/j.neuroimage.2006.02.024

Smith, S. M., Jenkinson, M., Woolrich, M. W., Beckmann, C. F., Behrens, T. E., Johansen-Berg, H., … Matthews, P. M. (2004). Advances in functional and structural MR image analysis and implementation as FSL. Neuroimage, 23 *Suppl 1*, S208–219. doi:10.1016/j.neuroimage.2004.07.051

Smith, S. M., & Nichols, T. E. (2009). Threshold-free cluster enhancement: Addressing problems of smoothing, threshold dependence and localisation in cluster inference. Neuroimage, 44(1), 83–98. doi:10.1016/j.neuroimage.2008.03.061

Soares, J. M., Marques, P., Alves, V., & Sousa, N. (2013). A hitchhiker’s guide to diffusion tensor imaging. Frontiers in Neuroscience, 7, 1–14. doi:10.3389/fnins.2013.00031

Squire, L. R., & Zola-Morgan, S. (1991). The medial temporal lobe memory system. Science, 253(5026), 1380–1386. doi:10.1126/science.1896849

Sreedharan, R. M., Menon, A. C., James, J. S., Kesavadas, C., & Thomas, S. V. (2015). Arcuate fasciculus laterality by diffusion tensor imaging correlates with language laterality by functional MRI in preadolescent children. Neuroradiology, 57(3), 291–297. doi:10.1007/s00234-014-1469-1

Su, M., Thiebaut de Schotten, M., Zhao, J., Song, S., Zhou, W., Gong, G., … Shu, H. (2018). Vocabulary growth rate from preschool to school-age years is reflected in the connectivity of the arcuate fasciculus in 14-year-old children. Developmental Science, 21(5), e12647. doi:10.1111/desc.12647

Sugiura, L., Ojima, S., Matsuba-Kurita, H., Dan, I., Tsuzuki, D., Katura, T., & Hagiwara, H. (2011). Sound to language: different cortical processing for first and second languages in elementary school children as revealed by a large-scale study using fNIRS. Cerebral Cortex, 21(10), 2374–2393. doi:10.1093/cercor/bhr023

Szaflarski, J. P., Schmithorst, V. J., Altaye, M., Byars, A. W., Ret, J., Plante, E., & Holland, S. K. (2006). A longitudinal functional magnetic resonance imaging study of language development in children 5 to 11 years old. Annals of Neurology, 59(5), 796–807. doi:10.1002/ana.20817

Takashima, A., Bakker-Marshall, I., van Hell, J. G., McQueen, J. M., & Janzen, G. (2019). Neural correlates of word learning in children. Developmental Cognitive Neuroscience, 37, 100649. doi:10.1016/j.dcn.2019.100649

Tamaki, M., Matsuoka, T., Nittono, H., & Hori, T. (2008). Fast sleep spindle (13–15 Hz) activity correlates with sleep-dependent improvement in visuomotor performance. SLEEP, 31(2), 204–211.

Tamminen, J., Lambon Ralph, M. A., & Lewis, P. A. (2013). The role of sleep spindles and slow-wave activity in integrating new information in semantic memory. Journal of Neuroscience, 33(39), 15376–15381. doi:10.1523/JNEUROSCI.5093-12.2013

Tamminen, J., Lambon Ralph, M. A., & Lewis, P. A. (2017). Targeted memory reactivation of newly learned words during sleep triggers REM-mediated integration of new memories and existing knowledge. Neurobiology of Learning and Memory, 137, 77–82. doi:10.1016/j.nlm.2016.11.012

Tamminen, J., Payne, J. D., Stickgold, R., Wamsley, E. J., & Gaskell, M. G. (2010). Sleep spindle activity is associated with the integration of new memories and existing knowledge. Journal of Neuroscience, 30(43), 14356–14360. doi:10.1523/JNEUROSCI.3028-10.2010

Tsanas, A., & Clifford, G. D. (2015). Stage-independent, single lead EEG sleep spindle detection using the continuous wavelet transform and local weighted smoothing. Frontiers in Human Neuroscience, 9, 181. doi:10.3389/fnhum.2015.00181

Torres, A. R., Mota, N. B., Adamy, N., Naschold, A., Lima, T. Z., Copelli, M., Weissheimer, J., Pegado, F., & Ribeiro, S. (2021). Selective Inhibition of Mirror Invariance for Letters Consolidated by Sleep Doubles Reading Fluency. Current Biology: CB, 31(4), 742– 752.e8.

Tucker, M. A., & Fishbein, W. (2008). Enhancement of declarative memory performance following a daytime nap is contingent on strength of initial task acquisition. SLEEP 31(2), 197–203.

Urger, S. E., De Bellis, M. D., Hooper, S. R., Woolley, D. P., Chen, S. D., & Provenzale, J. (2015). The superior longitudinal fasciculus in typically developing children and adolescents: Diffusion tensor imaging and neuropsychological correlates. Journal of Child Neurology, 30(1), 9–20. doi:10.1177/0883073813520503

van Schalkwijk, F. J., Sauter, C., Hoedlmoser, K., Heib, D. P. J., Klosch, G., Moser, D., … Schabus, M. (2019). The effect of daytime napping and full-night sleep on the consolidation of declarative and procedural information. Journal of Sleep Research, 28(1), e12649. doi:10.1111/jsr.12649

Vien, C., Bore, A., Boutin, A., Pinsard, B., Carrier, J., Doyon, J., & Fogel, S. (2019). Thalamo-cortical white matter underlies motor memory consolidation via modulation of sleep spindles in young and olderadults. Neuroscience, 402, 104–115. doi:10.1016/j.neuroscience.2018.12.049

Von Der Heide, R. J., Skipper, L. M., & Olson, I. R. (2013). Anterior temporal face patches: a meta-analysis and empirical study. Frontiers in Human Neuroscience, 7, 17. doi:10.3389/fnhum.2013.00017

Vukovic, N., Hansen, B., Lund, T. E., Jespersen, S., & Shtyrov, Y. (2021). Rapid microstructural plasticity in the cortical semantic network following a short language learning session. PLoS Biology, 19(6), e3001290. doi:10.1371/journal.pbio.3001290

Walker, M. P., & Stickgold, R. (2010). Overnight alchemy: sleep-dependent memory evolution. Nature Reviews Neuroscience, 11(3), 218; author reply 218. doi:10.1038/nrn2762-c1

Weighall, A. R., Henderson, L. M., Barr, D. J., Cairney, S. A., & Gaskell, M. G. (2017). Eye-tracking the time-course of novel word learning and lexical competition in adults and children. Brain & Language, 167, 13–27. doi:10.1016/j.bandl.2016.07.010

Wen, H., & Liu, Z. (2016). Separating Fractal and Oscillatory Components in the Power Spectrum of Neurophysiological Signal. Brain Topography, 29(1), 13–26. doi:10.1007/s10548-015-0448-0

Werchan, D. M., Kim, J. S., & Gomez, R. L. (2021). A daytime nap combined with nighttime sleep promotes learning in toddlers. Journal of Experimental Child Psychology, 202, 105006. doi:10.1016/j.jecp.2020.105006

Wickham, H. (2016). ggplot2: Elegant Graphics for Data Analysis. New York: Springer-Verlag.

Wilhelm, I., Prehn-Kristensen, A., & Born, J. (2012). Sleep-dependent memory consolidation - What can be learnt from children? Neuroscience and Biobehavioral Reviews, 36(7), 1718–1728. doi:10.1016/j.neubiorev.2012.03.002

Wilhelm, I., Rose, M., Imhof, K. I., Rasch, B., Buchel, C., & Born, J. (2013). The sleeping child outplays the adult’s capacity to convert implicit into explicit knowledge. Nature Neuroscience, 16(4), 391–393. doi:10.1038/nn.3343

Wixted, J. T. (2004). The psychology and neuroscience of forgetting. Annual Review of Psychology, 55, 235–269. doi:10.1146/annurev.psych.55.090902.141555

Zhang, Z. Y., Campbell, I. G., Dhayagude, P., Espino, H. C., & Feinberg, I. (2021). Longitudinal analysis of sleep spindle maturation from childhood through late adolescence. Journal of Neuroscience, 41(19), 4253–4261. doi:10.1523/JNEUROSCI.2370-20.2021

